# Arabidopsis SWEET12 regulates sugar allocation and defense responses to sustain beneficial association with *Serendipita indica* in roots

**DOI:** 10.1101/2025.09.29.679152

**Authors:** Abhimanyu Jogawat, Madhan Sanyasi, Sidhardh H. Menon, Divya Goyal, Athira Mohandas Nair, Jyothilakshmi Vadassery

## Abstract

Carbon availability is a central determinant of beneficial plant–fungal associations, and sugar transporters are key levers of this exchange. SWEETs (SUGARS WILL EVENTUALLY BE EXPORTED TRANSPORTER) are involved in transporting various kinds of sugars in plants; however, their functional roles in fungal symbiosis are not sufficiently explored. In this study, we investigate the functional relevance of Arabidopsis SWEETs in the interaction with endophytic fungi, *Serendipita indica*. Transcript profiling of SWEET genes in response to *S. indica* and its major elicitor, cellotriose, revealed early root-specific induction of SWEET12. Using a *SWEET12* loss-of-function mutant, we demonstrate that the absence of SWEET12 disrupts the major outcomes of mutualism including growth promotion, balanced colonization, sugar allocation, and the accumulation of defense phytohormones (JA and SA). Transcriptome profiling further reveals that SWEET12 buffers whole-plant responses by coordinating genes linked to carbohydrate, nitrogen, and lipid metabolism, and by tuning defense signalling and nutrient transporter networks. Our findings indicate that SWEET12 is essential for balancing fungal colonization and host defense, thereby promoting plant growth. SWEET12 does so by acting as sugar valve that meters sugar release to the apoplast, enabling the fungus to access carbon while preserving host sugar homeostasis and immune competence.

## 1. Introduction

Arbuscular mycorrhiza (AM) contribute an estimated 3.93 gigatons CO_2_ equivalents annually, while ectomycorrhiza and ericoid mycorrhiza contribute 9.07 and 0.12 gigatons, respectively underscoring their contribution in carbon sequestration (Hawkins et al. 2023). Carbon exchange and translocation in the form of sugars is not just a metabolic transaction but is a cornerstone of plant growth, development, and defense responses to biotic and abiotic stressors (Smith and Stitt 2007, Anjali et al. 2020). The importance of sugar transport in symbiosis becomes even clearer when considering the carbon economy of plants. During symbiosis, the fungal partner relies heavily on sugars as its primary carbon source, highlighting the critical role of sugar transporters in these interactions. Plants employ a diverse array of sugar transporters, including sucrose transporters (SUCs or SUTs), monosaccharide transporters (MSTs), and the recently discovered SWEET family. These transporters, characterized by their transmembrane domains and proton-coupled transport mechanisms, are pivotal in regulating sugar flow within plants (Doidy et al. 2012, Julius et al. 2017, Liu et al. 2025). Plant utilizes carbon source as a currency while trading for nutrient with symbiotic microorganism, but this trade is disrupted when pathogens interact with plants (Fellbaum et al. 2012, Rani et al. 2021, Chen et al. 2024). During the vegetative stage, plants are usually composed of one source (mature leaves) and two sinks (developing leaves and roots) (Durand et al. 2016). Roots are thus typical sink organs and depend exclusively on sucrose imported from the source leaves.

The SWEET (SUGARS WILL EVENTUALLY BE EXPORTED TRANSPORTER) family, in particular, has emerged as a key player in plant-microbe interactions. These bidirectional sugar efflux transporters, with their unique seven-transmembrane domain structure, are not only essential for plant development such as nectar secretion and seed formation, but also play a dual role in pathogen interactions (Chen et al. 2010, Baker et al. 2012, Zheng et al. 2024). Plant pathogens, both bacterial and fungal, hijack SWEET transporters to redirect sugar flow toward infection sites, ensuring their survival and proliferation, for instance, Arabidopsis SWEET2 by *Pythium* and *Trichoderma harzianum* (Chen et al. 2015, Rouina et al. 2021), grapevine *Vv*SWEET4 and *Vv*SWEET7 by *Botrytis cinerea* (Chong et al. 2014, Breia et al. 2020), cotton *Gh*SWEET42 by *Verticillium dahliae* (Sun et al. 2021), sweet potato *Ib*SWEET10 by *Fusarium oxysporum* (Li et al. 2017), potato *St*SWEET10c and *St*SWEET11 by *Phytophthora infestans* (Wu et al. 2024), *At*SWEET11 by *Pseudomonas syringae* (Fatima and Senthil-Kumar 2021), wheat *Ta*SWEET14d by stripe rust fungus (Zheng et al. 2025), citrus SWEET15 by *Xanthomonas citri* pv. c*itri* (Khadgi et al. 2025), tomato *Sl*SWEET17 by root-knot nematode (Wang et al. 2025), and soybean *Gm*SWEET20 by cyst nematode (Wu et al. 2025). The induction of host SWEET family by TAL (transcriptional activator like)-effectors is crucial for virulence of the bacterial rice pathogen *Xanthomonas oryzae pv. oryzae* (Xoo) (Chen et al. 2010). Four different bacterial TAL effectors are known to induce the rice sugar transporters *Os*SWEET11 and *Os*SWEET14 (Chu et al. 2006, Yang et al. 2006, Antony et al. 2010, Chen et al. 2010, Yuan et al. 2011). *Plasmodiophora brassicae* infection in Arabidopsis triggers the accumulation of SWEET11 and SWEET12 in phloem tissues, facilitating sugar transport to the pathogen in shoot and root (Walerowski et al. 2018). *Colletotrichum higginsianum* and *Botrytis cinerea* exploit SWEET transporters to enhance their virulence, with knockdowns of *SWEET11* and *SWEET12* conferring resistance against these pathogens (Chen et al. 2010, Gebauer et al. 2017).

When plants interact with beneficial microbes, the induction of SWEET sugar transporters in colonized roots plays a pivotal role in shaping these symbiotic relationships. In AM symbiosis with potato, for instance, vacuolar membrane-localized SWEETs like *SWEET2, SWEET16*, and *SWEET17*, along with plasma membrane-localized *SWEET11* and *SWEET12*, have been shown to be upregulated (Manck-Götzenberger and Requena 2016). In *Solanum tuberosum* and *Glycine max*, specific SWEET transporter genes have shown upregulation during mycorrhizal symbiosis (Zhao et al. 2019). SWEET14 mediates mycorrhizal symbiosis and carbon allocation in orchid, *Dendrobium officinale* (Li et al. 2025). Similarly, in Arabidopsis, SWEET11 and SWEET12 have shown to be involved in promoting plant growth by facilitating sucrose transport to a plant growth promoting rhizobia, *Pseudomonas simiae* WCS417r (Desrut et al. 2020). A recent study with Arabidopsis roots has further highlighted the importance of *SWEET2, SWEET4, SWEET11*, and *SWEET12* in maintaining beneficial root microbiota along the root axis in the rhizosphere by secreting sugars (Loo et al. 2024). The root endosymbiont, *Serendipita indica* stands out as a versatile model for studying plant-fungal mutualism. Unlike AMF, *S. indica* is axenically cultivable, and exhibits no host specificity, making it an ideal candidate for exploring symbiotic interactions across a wide range of plants including Arabidopsis (Verma et al. 1998, Jogawat et al. 2013, 2016, 2020, Kundu et al. 2022). Despite these advances, the role of SWEET transporters in *Serendipita indica* symbiosis remains underexplored. During root colonization, various factors of plant defense such as calcium signaling, ROS, callose deposition, glucosinolates and phytohormonal defense are crucial for maintaining and sustaining this beneficial interaction (Jogawat et al. 2020, Kundu and Vadassery 2022). One of the key features of *S. indica* is its ability to acquire sugars from host plants via fungal hexose transporters (Rani et al. 2015, Raj et al. 2021). While no plant sugar transporter has been specifically reported in *S. indica* interactions, recent RNA sequencing data revealed that transcripts of *SWEET3, SWEET11*, and *SWEET12* are upregulated in roots at 10 days post-inoculation (Pérez-Alonso et al. 2022). Intriguingly, Gandhi *et al*. (2024) demonstrated that *S. indica* releases a symbiotic signal, cellotriose (CT), which induces the expression of *SWEET11* and *SWEET12*. These findings suggest a regulatory network governing sugar transport during *S. indica* symbiosis. However, the precise functional roles of these SWEET transporters at different stages of symbiosis are unexplored.

In the present study, we show SWEET12 as an important sugar transporter during *S. indica* symbiosis. We delve into the functional role of SWEET12 during *S. indica* colonization in Arabidopsis roots, a phase where sugar transport becomes pivotal for balanced colonization and sustaining the fully established mutualistic relationship. Our findings reveal that SWEET12 is a critical player in facilitating *S. indica* colonization and driving growth promotion via regulating sugar availability and affecting plant immunity.

## 2. Materials and methods

### 2.1. Plant and Fungal Material and Growth Conditions

The root endosymbiont *Serendipita indica* (syn. *Piriformospora indica*; Verma et al. 1998) was cultured and maintained on modified Kaefer’s (KF) medium at 28 ± 2°C with constant shaking at 110-150 rpm in an Infors HT multitron standard incubator shaker (Hill and Kafer 2001). For co-cultivation experiments on solid media and soil, wild-type *Arabidopsis thaliana* (WT; ecotype Columbia-0) and *sweet12* (At5g23660; SALK_031696.56.00.x) mutant line were used (Alonso et al. 2003). This mutant line was earlier reported to be knock-out line (Chen et al. 2012) and have been used in previous studies (Desrut et al. 2020, Fatima et al. 2021). Additionally, we also confirmed the reduced level of SWEET12 transcripts by real time PCR (**Fig. S1**). Seedlings and adult plants were grown in a growth room at the set long day parameters i.e. temperature 22°C, a photoperiod of 14h light/10h dark, and a light intensity of 150–170 µmol per square meter per second (m^-2^ s^-1^).

### 2.2. Co-cultivation of S. indica and Arabidopsis in Media and Soil Conditions

For co-cultivation on solid media, surface-sterilized *Arabidopsis* seeds were stratified and germinated on half-strength Murashige and Skoog (MS) medium for 7 days at 22°C under a 14h light/10h dark photoperiod, with a light intensity of 150 µmol m^-2^ s^-1^ in a growth chamber (Percival). Co-cultivation with *S. indica* was carried out following the protocol established in our previous study (Johnson et al. 2011, Jogawat et al. 2020). In brief, seven-day-old seedlings were transferred to modified 1× Poor Nutrient Medium (PNM) plates, with borer-cut *S. indica*-KF agar plugs placed between them and samples were harvested at 7, 14, and 21 days post-inoculation (dpi) for further analysis. For soil-based co-cultivation, stratified seeds were first grown in horticulture-grade soilrite, a mixture of Irish peat moss, perlite, and exfoliated vermiculite in a 1:1:1 ratio for 2 weeks. These plants then transferred in another pots containing the soil properly pre-mixed with 1% (w/w) *S. indica* mycelia to ensure consistent colonization. Plants were maintained under the same growth conditions (22°C, 14 h light/10 h dark photoperiod, 150 µmol m^-2^ s^-1^ light intensity for 4 weeks. Samples were collected at 30 dpi for further analysis.

### 2.3. Treatments and expression profiling of SWEETs

Stratified seeds were grown for 7 days on MS plates, after which the seedlings were exposed to a 10 µM cellotriose (Sigma-Aldrich, C1167) treatment supplemented in liquid ½-strength MS medium. Samples were harvested at 0 min, 30 min, 1 hr, 2 hrs, 4 hrs, and 8 hrs. Roots from co-cultivated samples were harvested at 7 and 14 dpi. After freezing the samples in liquid N_2_, they were ground into a fine powder along with the TRIzol Reagent (Invitrogen) and total RNA was extracted following the manufacturer’s instructions. Further, DNAse treatment (Turbo DNAse, Ambion) was performed to eliminate any DNA contamination. Finally, cDNA was synthesized using the High-Capacity cDNA Kit (Applied Biosystems, Thermo Fisher Scientific). For Real-Time PCR, gene-specific primers were designed using the NCBI primer design tool (http://www.ncbi.nlm.nih.gov/tools/primer-blast). Quantitative RT-PCR was conducted in optical 96-well plates on a CFX96 Real-Time PCR Detection System (Bio-Rad), using 6 μL of iTaq Universal SYBR Green Mix (Bio-Rad), 2 μL of 4× diluted cDNA, and gene-specific forward and reverse primers. The endogenous control *AtActin2* (*At3g18780*) was used for normalization of transcripts. Fold induction values of target genes were calculated using the ΔΔCT equation (Livak and Schmittgen 2001), with the mRNA level of target genes in non-treated seedlings set to 1.0. The primer pairs used are listed in **Supplementary Table S1**.

### 2.4. Estimation and detection of S. indica colonization

*S. indica*-colonized roots were treated with fungal specific Alexa Flour 488-WGA fluorescent dye to stain mycelia as previously described by Kaladhar et al. (2023), and the roots were observed under Laser Scanning Microscope (AOBS TCS-SP8; LEICA GERMANY) at an emission wavelength of 505–530 nm and excitation at 470 nm, with optical sectioning at a step size of 1µm across the root. To quantify the depth of penetration, the optically sectioned images were stacked into z-plane to generate a 3D image with scales of each image representing the depth penetrated by the fungus. The relative fungal DNA amount was also quantified using *Arabidopsis Actin2* (At3g18780) and *S. indica Tef1* primers in real-time PCR (Bütehorn et al. 2000). Fold changes in relative fungal DNA amount were calculated using the CT of *SiTef1* normalized by the CT of *AtActin2* with the ΔΔCT equation, and the *S. indica* DNA content in control roots was set to 1.0 (Vadassery et al. 2008, Jogawat et al. 2020).

### 2.5. Tissue localization by β-glucosidase (GUS) assay

Transgenic Arabidopsis lines expressing the *SWEET12* promoter fused to β-glucuronidase (*ProSWEET12*:*GUS*), previously developed by Le Hir et al. (2015), was used in this study. These T3-generation seedlings were treated with 10 µM CT in 1 mL of half-strength MS medium. These seedlings were also co-cultivated with *S. indica*. At 7 dpi, and 14 dpi, they were harvested, treated with GUS staining buffer (0.1% Triton X-100, 50 mM NaHPO_4_ buffer (pH 7.2), 2 mM potassium Ferrocyanide, 2 mM potassium Ferricyanide, and 1 mM 5-bromo-chloro-3-glucuronide), and incubated in the dark at 37°C for 12 hours (Li 2011). The tissues were treated with a freshly prepared decolorizing solution of ethanol:50% glycerol:acetone (9:3:3) at room temperature and finally observed under a light microscope (Stereozoom Microscope, Nikon).

### 2.6. Soluble sugars levels analysis by targeted GC-MS

The sugar estimation method by Kundu et al. 2022 was used. In brief, *S. indica* colonized samples at 30 dpi were collected, lyophilized, and ground into a fine powder. From this, 20-25 mg of powder was used to extract sugars with 480 µL of 100% methanol, and 20 µL of 0.2 mg/mL ribitol (adonitol) was added as an internal standard. The mixture was vigorously shaken for 2 minutes, then heated at 70°C for 15 minutes. After that, an equal volume of water was added, and the mixture was shaken again, followed by the addition of 250 µL of chloroform and thorough mixing. The mixture was centrifuged at 2200 g for 10 minutes at room temperature. The upper aqueous phase was removed and dried in a speed vacuum rotator at 35°C. The dried fraction was then redissolved in 40 µL of 20 mg/mL methoxamine hydrochloride in pyridine and incubated for 90 minutes at 37°C. Afterward, 60 µL of MSTFA (N-methyl-N-(trimethylsilyl) trifluoroacetamide) was added, and the mixture was incubated for 30 minutes at 37°C. After the derivatization process, the sample was transferred to a gas chromatography–mass spectrometry (GC-MS) vial containing an insert. The injection volume was set at 0.2 µL, and the injection mode was set to Split Mode with a split ratio of 5. GC-MS analysis for targeted sugar analysis was conducted using a Shimadzu Gas Chromatograph (GC-2010 Plus) coupled with a mass spectrometer (TQ 8050) and an auto sampler (AOC-20s)–auto injector (AOC-20i). Analysis was performed with a SH-Rxi-5Sil MS capillary column (30 m x 0.25 µm, 0.25 mm) (Restek Corporation, USA), and helium was used as the carrier gas with a flow rate of 1 mL/min. The method consisted of an 80°C isothermal heating for 2 minutes, followed by a ramp rate of 5°C/min to 250°C, with a 2-minute hold, and a final ramp of 10°C/min with a 24 minutes hold. The total run time for GC-MS was 67 minutes with a solvent delay of 4.5 minutes. Chromatogram integration and mass spectrum analysis were performed using Shimadzu Lab Solutions software (GC-MS Solution Version 4.53SP1), and the NIST17 spectral library was used for derivatized metabolite identification.

### 2.7. Phytohormone estimation

Defense phytohormones were quantified as described in our previous studies (Vadassery et al. 2012). In brief, samples from control and *S. indica*-colonized seedlings at 7 dpi were harvested, immediately frozen in liquid N_2_, lyophilized, and ground into a fine powder. For phytohormone quantification, each pre-weighed sample (25 mg) was extracted using 1 mL of methanol containing 40 ng of d6-jasmonic acid (HPC Standards GmbH, Cunnersdorf, Germany), 40 ng of salicylic acid-d4 (Santa Cruz Biotechnology, Santa Cruz, U.S.A.), 40 ng of abscisic acid-d6 (Toronto Research Chemicals, Toronto, ON, Canada) as internal standards. Phytohormones were quantified on Exion LC coupled to Triple Quad 6500+ (Sciex). Samples were loaded onto Zorbax Eclipse XDB-C18 (50x4.6mm, 1.8um, Agilent Technologies) column with flow rate of 1.1ml/min. Scheduled Multiple-reaction monitoring (MRM) was used to monitor analyte parent ion → product ion with detection window of 60sec. Phytohormones were quantified relative to the signal of their corresponding internal standard

### 2.8. Total RNA isolation, library preparation and RNA-sequencing analysis

To analyse differentially expressed genes upon *S. indica* colonization with WT and *sweet12*, total RNA was isolated from pooled plant roots (∼1500 roots / sample; 1500*3 = 4500) as described earlier (Jogawat et al. 2020). The quantity of total RNA was assessed by using a Nanodrop® ND-1000 spectrophotometer (Thermo Fisher Scientific). Further, RNA integrity and quality was also checked on a Bioanalyzer 2100 (Agilent). For whole transcriptome analysis, the RNA samples were used for cDNA library preparation (PE150), RNA sequencing was done on Illumina NovaSeq™ 6000 platforms. The obtained raw fastq reads were processed using Fastp v.0.23.4 (Chen et al. 2018) and thereafter detection and removal of rRNA sequences was performed using RiboDetector v.0.3.1 (Deng et al. 2022). The filtered reads were aligned to STAR indexed *Arabidopsis thaliana* sequence (Ensembl, TAIR10) using STAR aligner v2.7.9a (Dobin et al. 2013). The rRNA and tRNA features were removed from the GTF file of the *Arabidopsis thaliana*. The alignment file (sorted BAM) from individual samples were quantified using FeatureCounts v.2.0.1 (Liao et al. 2014) based on the filtered GTF file, to obtain gene counts. Differential expression estimation was done based on the gene counts using DESeq2 at p-value 0.05 (Love et al. 2014). The annotated DESeq2 result file was filtered based on Adjusted p-value (FDR) ≤ 0.05 and Log Fold Change ±0.58.

### 2.9. Bioinformatics analyses

The significant genes from the different groups were subjected to GO and KEGG enrichment analysis using ShinyGO (Ge et al. 2020) and AgriGo v2.0 (Tian et al. 2017) with the *Arabidopsis thaliana* genome as the reference model, a significance threshold of 0.05 and False Discovery Rate (FDR) as the adjustment method. Further, the LogFC ±0.58 (equal to fold change 1.5) was used for identifying differentially expressed genes (DEGs), dot plot generation and DiVenn diagrams (Ge et al. 2020, Sun et al. 2019). For clustering analysis, we used iDEP2.0 tool (https://bioinformatics.sdstate.edu/idep/) for k-means clustering managed by South Dakota State University, USA with FDR cut-off (0.05) (Ge et al. 2018). Dot plots representing comparative ontologies were generated using R.

### 2.10. Statistical analysis

We used Student’s *t*-test or one- or two-way ANOVA with Tukey’s test in Prism software to statistically analyze control and treated data. Asterisks or different letters were used to indicate significance among different treatments and genotypes. Figures were generated using Graph Pad Prism version 8.4.2.

## 3. Results

### 3.1. Serendipita indica colonization enhances sugar levels in host plants

To investigate the effect of sugars on *S. indica* symbiosis, we quantified the levels of various sugars in both control and *S. indica*-colonized seedlings. At 21 days post-inoculation (dpi), *S. indica* colonization led to a significant increase in the levels of nearly all measured sugars, including key sugars such as sucrose, glucose, fructose, and galactose (**Fig. 1A and B**). This highlights the impact of *S. indica* on the host plant’s sugar metabolism during symbiosis. To further explore the influence of external sugar availability on this symbiotic relationship, we supplemented the co-cultivation PNM media with 1% sucrose and assessed its effects on plant growth promotion. Intriguingly, the addition of sucrose was sufficient to induce growth promotion, similar to *S. indica* and co-supplementation of both sugar and fungus did not confer any additional growth benefits under sucrose-supplemented conditions (**Fig. 1C**). This suggests that an external carbon source is crucial and excess can disrupt the mutualistic relationship between *S. indica* and its host plant. Controlled colonization on plant roots is crucial for *S. indica* symbiosis (Jogawat et al. 2020). In the presence of 1% sucrose, *S. indica* exhibited over-colonization at 7 and 14 dpi compared to control untreated plants (**Fig. 1D**). This behavior underscores the delicate balance of carbon exchange required to maintain a functional symbiosis

**Fig. 1.**
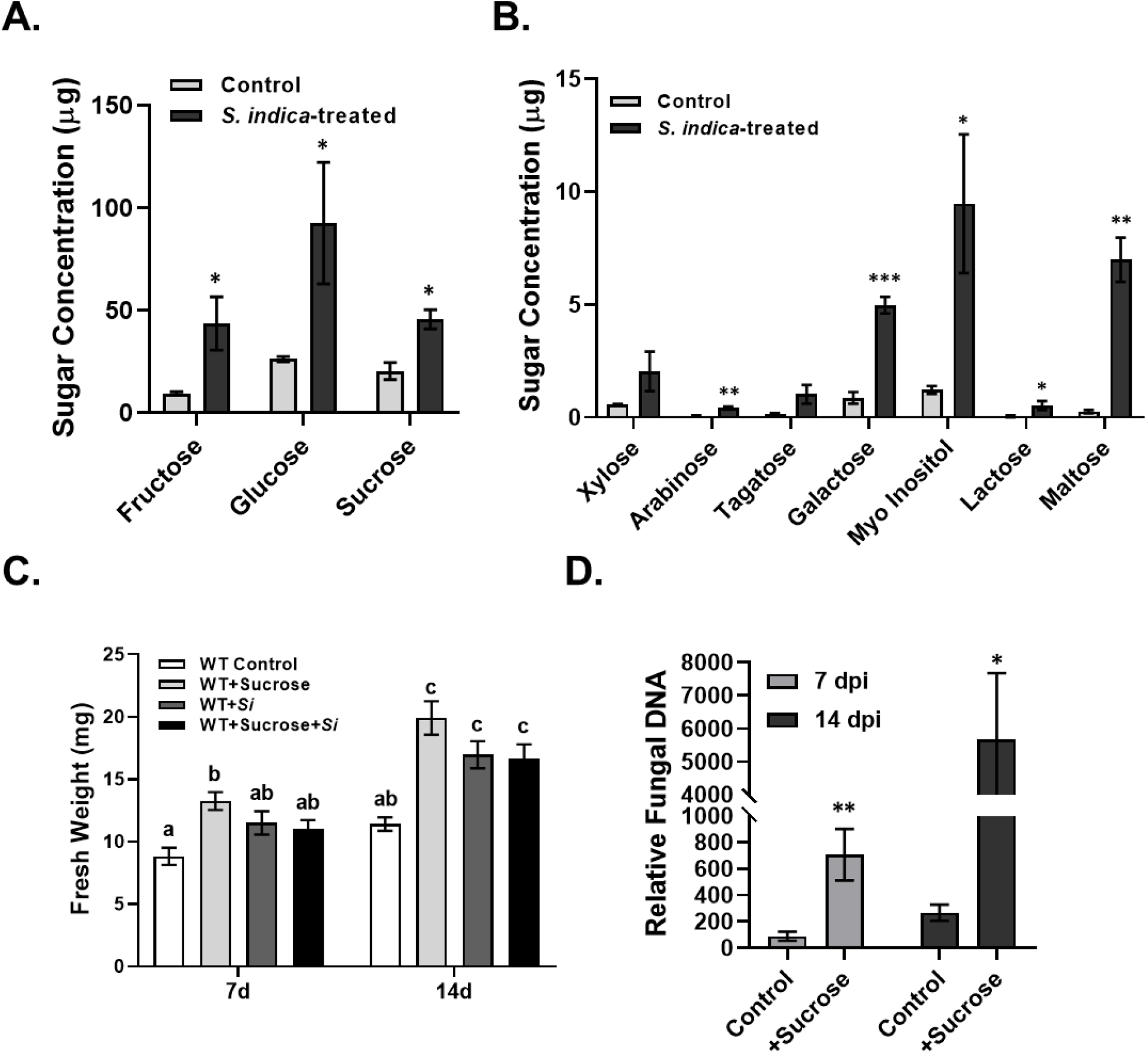
*S. indica-*induced sugar levels and growth promotion assay with exogenously added sucrose. **A, B. Sugar analysis during colonization:** At 21 dpi of *S. indica* colonization in Arabidopsis, sugar levels were measured using GC-MS/MS analysis. Data represents mean ± SEM where n= 3 (3*100=300). Asterisks indicate *p*-values as follows * = <0.05, ** = <0.005, and *** = <0.0005 after student’s *t*-test analysis. **C. Effect of sugar on *S. indica* symbiosis:** *S. indica-* mediated growth promotion was checked after co-cultivation with and without sucrose supply by measuring fresh weight. Data represents mean ± SEM of n= 20. Here, different letter shows significance difference after ANOVA analysis (*p* < 0.05, one-way ANOVA with Tukey’s test). **D. Relative Fungal DNA amount:** Arabidopsis was co-cultivated with *S. indica* on 1X PNM agar plates with or without 1% sucrose. Roots were harvested at 7 and 14 dpi and total genomic DNA were isolated. Relative fungal DNA in harvested roots was and quantified using RT-PCR to show colonization level in WT and *sweet12* mutant plant roots. Asterisks indicate significant differences after sucrose treatment over its corresponding control (* = p < 0.05, ** = p < 0.005; Student’s *t*-test). Data represents mean ± SEM where n= 4 independent biological replicates of roots (4*12=48).

### 3.2. Cellotriose as well as S. indica colonization induce SWEET family transporters

One of the symbiotic signal released from the cell wall of *S. indica*, cellotriose (CT), is a sugar that plays a key role in mediating this plant-fungal interaction (Vadassery et al. 2009a, Johnson et al. 2018). SWEET transporters, may serve as potential mediators of CT signaling and to explore the relationship, we treated Arabidopsis seedling roots with 10 µM CT and performed expression profiling of all 17 *AtSWEET* genes at 0, 4, and 8 hours post-treatment. Five *AtSWEET*s were induced at both time points, with *SWEET12* showing the strongest expression at 4 hours, followed by *SWEET1, 6, 10*, and *17* (**Fig. 2A**). Similarly, during *S. indica* colonization, seven *AtSWEETs* were found to be induced. Among them, *SWEET12* exhibited peak expression as early as 7 dpi, *SWEET 11* at 14 dpi, followed by *SWEET1, 4, 7, 13, 14*, and *15* in roots (**Fig. 2B**). Notably, *SWEET1, 7, 12, 14, 15* and *17* were commonly upregulated under both CT treatment and *S. indica* colonization. These findings highlight the dynamic expression patterns of SWEET transporter transcripts during CT treatment and *S. indica* symbiosis, suggesting potential roles in mediating sugar signaling during the establishment of this beneficial interaction.

**Fig. 2.**
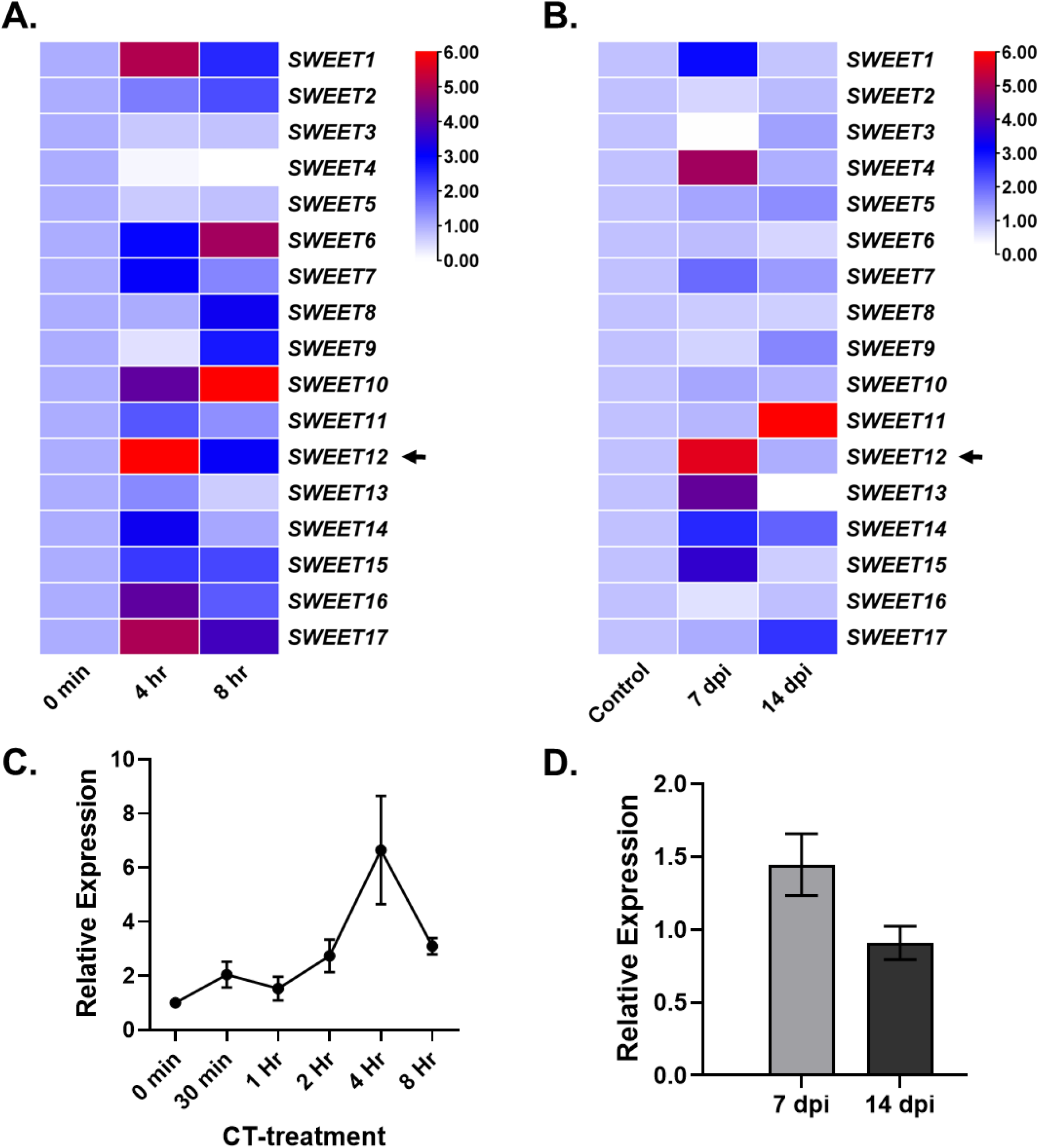
Expression analysis of *SWEET12* upon Cellotriose-treatment and *S. indica* colonization. **A. Expression analysis of *SWEETs* after CT-treatment:** Ten days old seedling roots were treated with 10 µM CT and the transcripts level of *SWEET* genes were checked after 0, 4 and 8 hr of CT-treatment. Data represents mean ± SEM of fold change (n=4*10=40). **B. Expression analysis of *SWEETs* in roots after *S. indica* treatment:** Seven days old seedlings were co-cultivated with *S. indica* on 1X PNM and the transcripts level of *SWEET* genes were checked after 7 days post inoculation (7 dpi) in roots. Data represents mean ± SEM of fold change (n = 4*12=48). **C. Time course expression analysis of *SWEET12* gene after CT-treatment:** Ten days old seedlings were treated with 10 µM CT and their expression was checked at 0, 30 min, 1 hr, 2 hr, 4 hr and 8 hr. Data represents mean ± SEM of fold change (n=4*10=40). **D. Expression of *SWEET12* in shoots at 7 and 14 dpi:** *SWEET* genes were checked after 7 dpi in shoots. Data represents mean fold change ± SEM of n = 3 (3*10=30).

### 3.3. SWEET12 shows early root specific expression on S. indica colonization

*SWEET12* was selected for further investigation based on its early induction and strongest upregulation in response to both CT-treatment and *S. indica* colonization (**Fig. 2A and B**). To assess the transcript kinetics of *SWEET12* in *S. indica* symbiosis, we analyzed the transcript levels of *SWEET12* at various time points i.e. 0, 30 minutes, 1 hr, 2 hr, 4 hr, and 8 hr. Our relative expression analysis revealed that *SWEET12* reached to its highest peak of expression at 4 hrs (**Fig. 2C**). Additionally, during *S. indica* colonization, *SWEET12* transcripts were induced at 7 dpi in *S. indica*-colonized roots (**Fig. 2B**), with minimal induction in the shoot (**Fig. 2D**). These results highlight the activation of *SWEET12* in during *Arabidopsis* and *S. indica* interaction. Further, to investigate the tissue specific expression patterns of *SWEET12 in-planta*, we utilized *ProSWEET12:GUS* reporter lines (Le Hir et al. 2015). Following treatment with 10 µM CT, *SWEET12* promoter activity was activated in roots, including the root vasculature, cortex, and root hairs [**Fig. 3A(a–f**)]. We also observed *SWEET12* promoter activity co-localization with *S. indica* spores in roots at 7 dpi. Similarly, during *S. indica* colonization, *SWEET12* promoter activity was primarily detected in the root vasculature, cortex, budding roots and root hairs roots at 7 dpi [**Fig. 3B(a–f**)]. These findings suggest that *S. indica* specifically regulates *SWEET12* expression in a root tissue-specific manner.

**Fig. 3.**
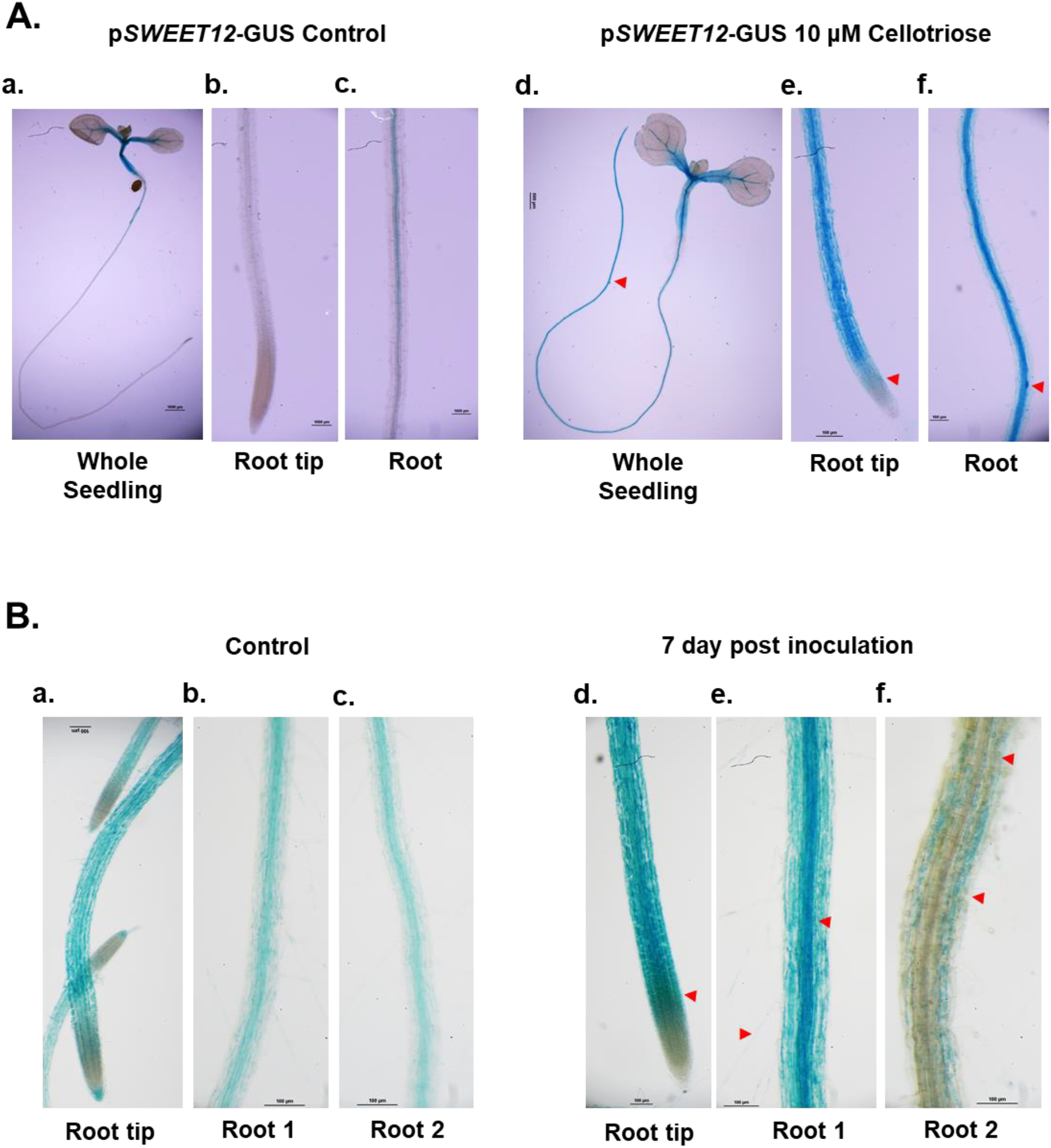
*SWEET12* promoter activity during *S. indica* symbiosis. **A. p*SWEET12*-*GUS* activity after CT-treatment:** Ten days old p*SWEET12*-*GUS* expressing seedlings were treated with 10 μM CT for 30 min and GUS assay was performed. Images of whole seedling (**a, d**), root tip (**b, e**), and root (**c, f**) were captured. The blue color indicates GUS expression induced by activity of *SWEET12* promoter in different tissues. Arrows indicate the induced GUS expression in different regions of root upon CT-treatment. **B. p*SWEET12*-*GUS* activity upon *S. indica* colonization:** p*SWEET12*-*GUS* expressing seedling were grown for 7 days on MS and then co-cultivated with *S. indica* on 1X PNM media. At 7 dpi, GUS assay was performed. The blue color shows *SWEET12* promoter activity at 7 dpi in different root parts. Images of root tip (**a, d**), root 1 with root hairs (**b, e**), root 2 (**c**) and root 2 with fungal spores (**f**) were captured. Arrows indicate the induced GUS expression in different regions of root upon *S. indica* colonization.

### 3.4. Loss-of-function of SWEET12 shows growth inhibition upon S. indica colonization

To explore the functional roles of SWEET12 in *S. indica*-mediated growth promotion, we co-cultivated *sweet12* mutant as well as wild-type (WT) seedlings with *S. indica*. At 14 dpi, *S. indica* colonization significantly enhanced total seedling growth in WT compared to untreated control, whereas the *sweet12* mutants showed growth inhibition upon *S. indica* treatment (**Fig. 4A and B**). This growth inhibition in *sweet12* mutant, was more pronounced in roots than in shoots (**Fig. 4C and D**). These results were further validated by soil-based co-cultivation experiments, where *sweet12* mutants exhibited growth inhibition, in contrast to the pronounced growth enhancement seen in WT plants at 30 dpi (**Fig. 4E and 4F**). These findings underscore the critical functional role of *SWEET12* in mediating *S. indica*-induced growth benefits.

**Fig. 4.**
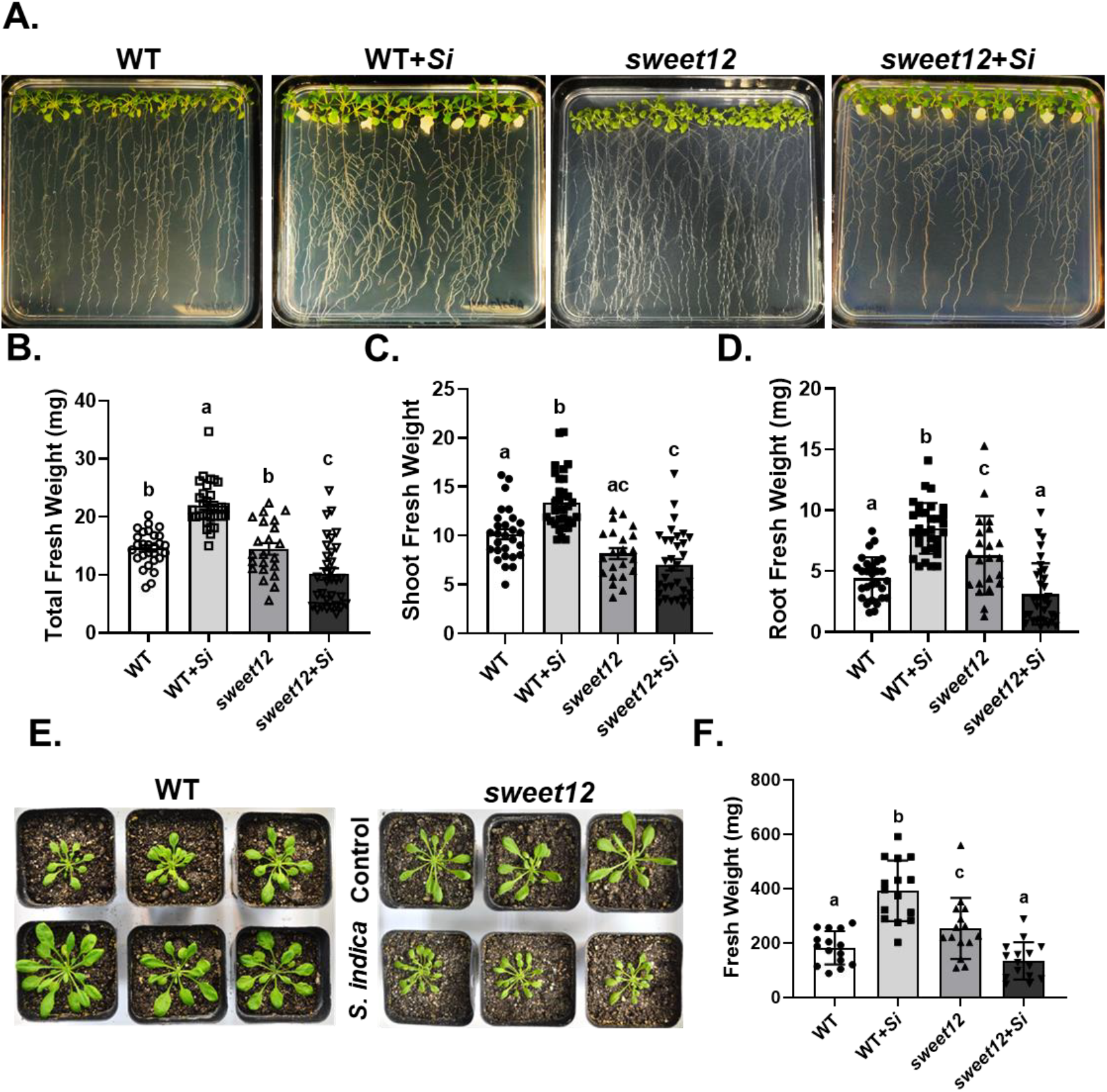
*S. indica* colonization with *sweet12* mutant. **A. Representative picture of co-cultivation on solid 1X PNM:** Seven days old WT and *sweet12* mutant were co-cultivated and growth promotion was observed at 14 dpi. **B. Total fresh weight, C. Shoot fresh weight, and D. Root fresh weight of WT and *sweet12* co-cultivated with *S. indica* on 1X PNM plates:** Seven days old seedlings were co-cultivated with *S. indica* and total fresh weight was measured. Data represents mean ± SEM with n= ≥22. Different letters indicate significance difference after ANOVA analysis (p < 0.05, one-way ANOVA with Tukey’s test). **E. Representative picture of soil co-cultivation:** Two weeks old Seedlings of WT and *sweet12* mutant were co-cultivated with 1% *S. indica* mixed in soil and the pictures were taken after one month (30 dpi). **F. Fresh weight of WT and *sweet12* co-cultivated with *S. indica* on soil:** After co-cultivation, rosette fresh weight was measured after one month of *S. indica* co-cultivation (30 dpi). Data represents mean ± SEM where n= 15. Different letters indicate significance difference after ANOVA analysis (p < 0.05, two-way ANOVA with Tukey’s multiple comparisons test.

### 3.5. S. indica colonization is altered in sweet12 mutant roots

To assess the cause of the growth inhibition in *SWEET12* loss-of-function mutant upon *S. indica* colonization, we quantified fungal DNA levels in *sweet12* mutant plant roots using a qPCR-based approach and compared with WT plant (Vadassery et al. 2012, Jogawat et al. 2020). The *S. indica*-specific gene, Translation elongation factor 1 (*SiTef1*) was used to estimate colonization levels, normalized against the *Arabidopsis Actin 2* (*AtAct2*) gene. Intriguingly, *sweet12* mutants exhibited significantly higher colonization levels than WT roots (**Fig. 5A**). Alexa Flour 488 WGA-stained fungal hyphae and mycelial network were observed mostly on the plant root surface with clumps and mycelial mass in *sweet12*, compared to lower fungal levels on WT roots at 7 and 14 dpi (**Fig. 5B**). To understand if increased fungal mycelia on root surface co-relates with increased penetration into roots, we analyzed the penetration levels and found that there was increased depth of fungal penetration in *sweet12* roots compared to WT at 7 and 14 dpi, as indicated by scale of the 3D images (**Fig. 5C)**. These results suggest that *SWEET12* play critical roles in regulating *S. indica* colonization on roots and are crucial for balanced colonization on roots.

**Fig. 5.**
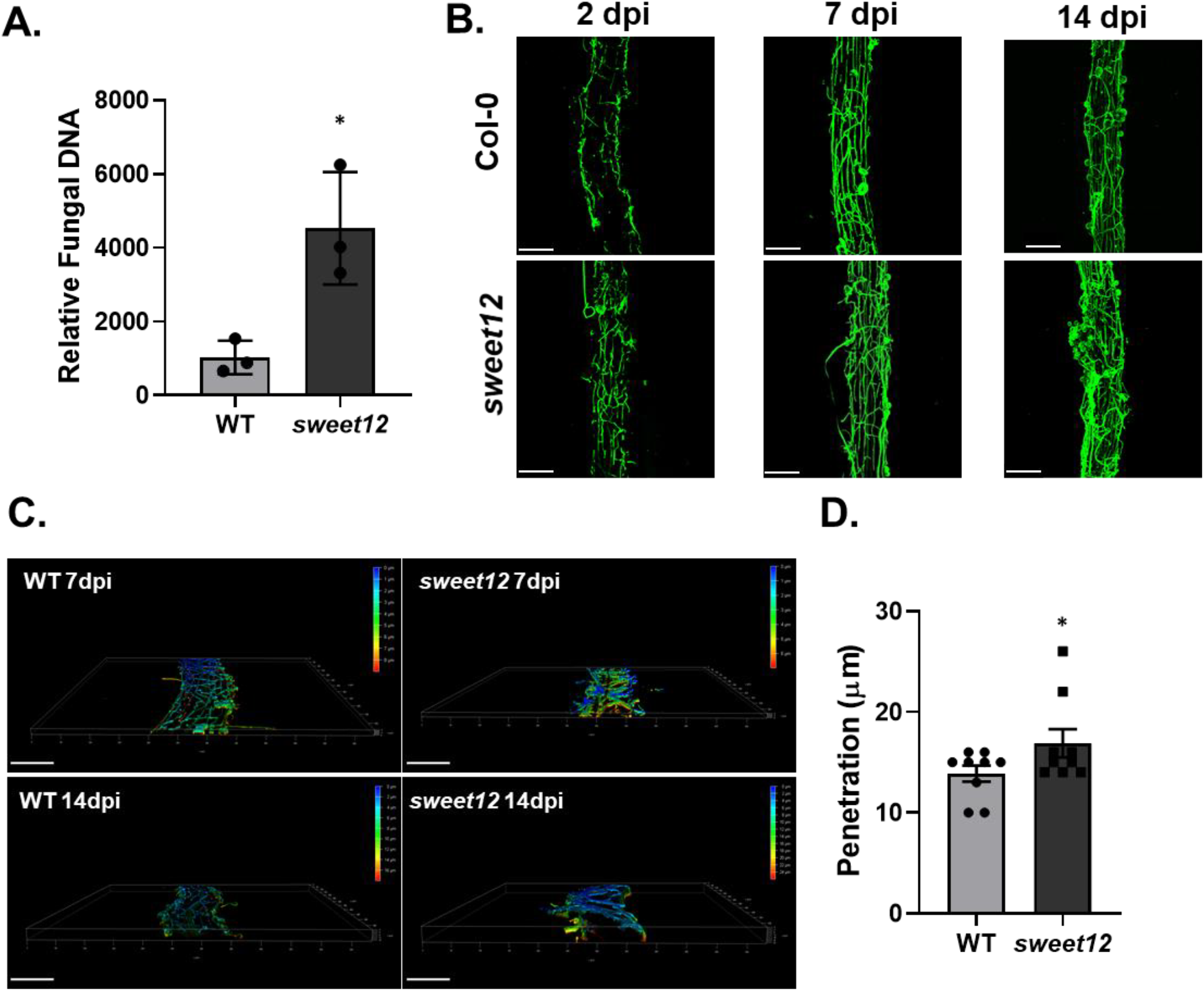
*S. indica* colonization and penetration of *sweet12* mutant. **A. Relative Fungal DNA amount:** Roots at 14 dpi were collected and genomic DNA was isolated for RT-PCR using *SiTef1*. Colonization levels were estimated in WT and *sweet12* mutant roots at 14 dpi. Asterisks indicate significant differences in *sweet12* mutant (p < 0.05, Student’s *t*-test). Data represents mean ± SEM of 3 independent biological replicates (3*12=36). **B. Representative picture of Alexa Flour 488-WGA stained S. *indica* in WT and *sweet12* mutant roots:** WT and *sweet12* mutant roots were harvested and stained with fungus staining florescent dye Alexa Flour 488-WGA and pictures were taken under confocal microscope. Scale bar = 100 μm. **B. Penetration Analysis of *S. indica* in plant roots:** Confocal images of *S. indica* colonization in wild-type (WT) and *sweet12* mutant roots were captured at 7 dpi (days post inoculation) using Alexa Fluor 488-labeled wheat germ agglutinin (WGA) staining. The images were optically sectioned and stacked into a Z-plane, illustrating the depth of fungal penetration into the roots. Imaging began at the surface of the root (0 μm) to track the presence of fungal colonization. The red color represents the deepest zone of fungal penetration, where *S. indica* mycelia were detected last, while green and blue indicate shallower regions, with blue representing the root surface (Scale bar = 100 μm). **D. Statistical analysis of *S. indica* penetration in roots:** The graph shows comparative depth of root tissue colonization by *S. indica* in WT and *sweet12* at 14 dpi. Asterisks indicate significant differences in *sweet12* mutant (p < 0.05, Student’s *t*-test). Data represents mean ± SEM of 10 plant roots.

### 3.6. The sweet12 mutants have altered sugar dynamics during S. indica colonization

Soluble sugars translocated to the roots can serve as a critical resource for *S. indica* colonization. Arabidopsis SWEET12 is capable of transporting glucose, fructose and sucrose (Le Hir et al. 2015). To investigate cause of growth inhibition by over-colonization in *sweet12* mutant roots, we estimated sugar level in *S. indica*-colonized plants. We quantified the levels of ten key sugars in whole plants using GC-MS/MS analysis. In wild-type (WT) plants, *S. indica* colonization significantly increased the levels of nearly all measured sugars at 21 and 30 dpi including sucrose, fructose and glucose. In *sweet12* mutant plants, there was constitutively high levels of sucrose, fructose, glucose *myo*-inositol and other sugars, which either did not change or decrease upon *S. indica* colonization (**Fig. 6A–C**). The enhanced sugar levels in *sweet12* mutant enhances *S. indica* colonization, which alters the balance required for efficient growth promotion.

**Fig. 6.**
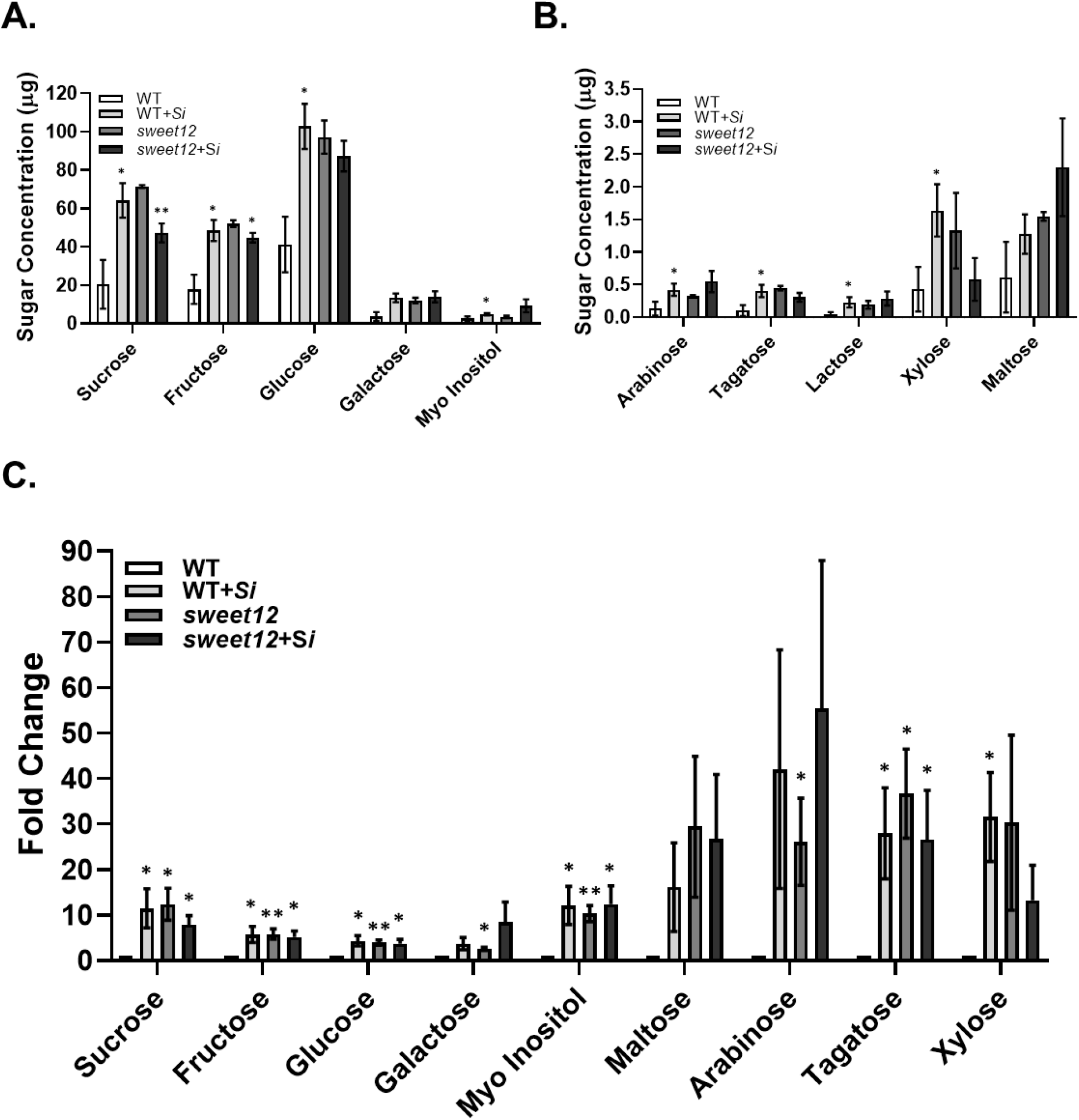
Sugar estimation by GC/MS analysis in WT and *sweet12*. **A, B. Sugar content in WT and *sweet12* mutant plants 30 dpi:** Different sugars were estimated in lyophilized and powdered plant material harvested at 30 dpi when symbiosis is fully established. Data represents mean ± SEM of 3 independent biological samples each having 4 rosettes. Asterisks indicate significant differences in *sweet12* mutant (*= P < 0.05, ** = p <0.005). **C. Fold change of sugar content in WT and *sweet12* mutant plants at 30 dpi:** Data are presented as mean ± SEM, calculated relative to the untreated WT control (set as 1-fold; n = 3*4 = 12). Asterisks indicate statistically significant differences compared to the untreated WT control, as determined by Student’s *t*-test: *p < 0.05, *p < 0.005.

### 3.7. Sugar imbalance affects defense phytohormones in S. indica-colonized sweet12 mutants

Given that soluble sugar levels are known to influence defense phytohormones such as JA and SA (Machado et al. 2016, Gebauer et al. 2017), we further analyzed the levels of these defense phytohormones in *S. indica*-colonized plants to explore potential links between sugar metabolism and defense signaling. To explore this interplay, we quantified the levels of key defense phytohormones JA and SA in control and *S. indica*-colonized seedlings. In WT seedlings, *S. indica* colonization significantly induced the levels of JA and these levels were further enhanced in *sweet12* mutant seedlings (**Fig. 7A**). We also observed an increase in SA levels in *S. indica*-colonized WT, but not in *sweet12* mutants (**Fig. 7B**). SA markers in *S. indica* symbiosis such as *CBP60g* and *PBS3* (Vahabi et al. 2015, Johnson et al. 2018, Pérez-Alonso et al. 2022) were not induced in *sweet12* roots, as in WT roots (**Fig. S4A**). These findings suggest that absence of *SWEET12* affects JA and SA levels potentially via influencing the defense-growth tradeoff.

**Fig. 7.**
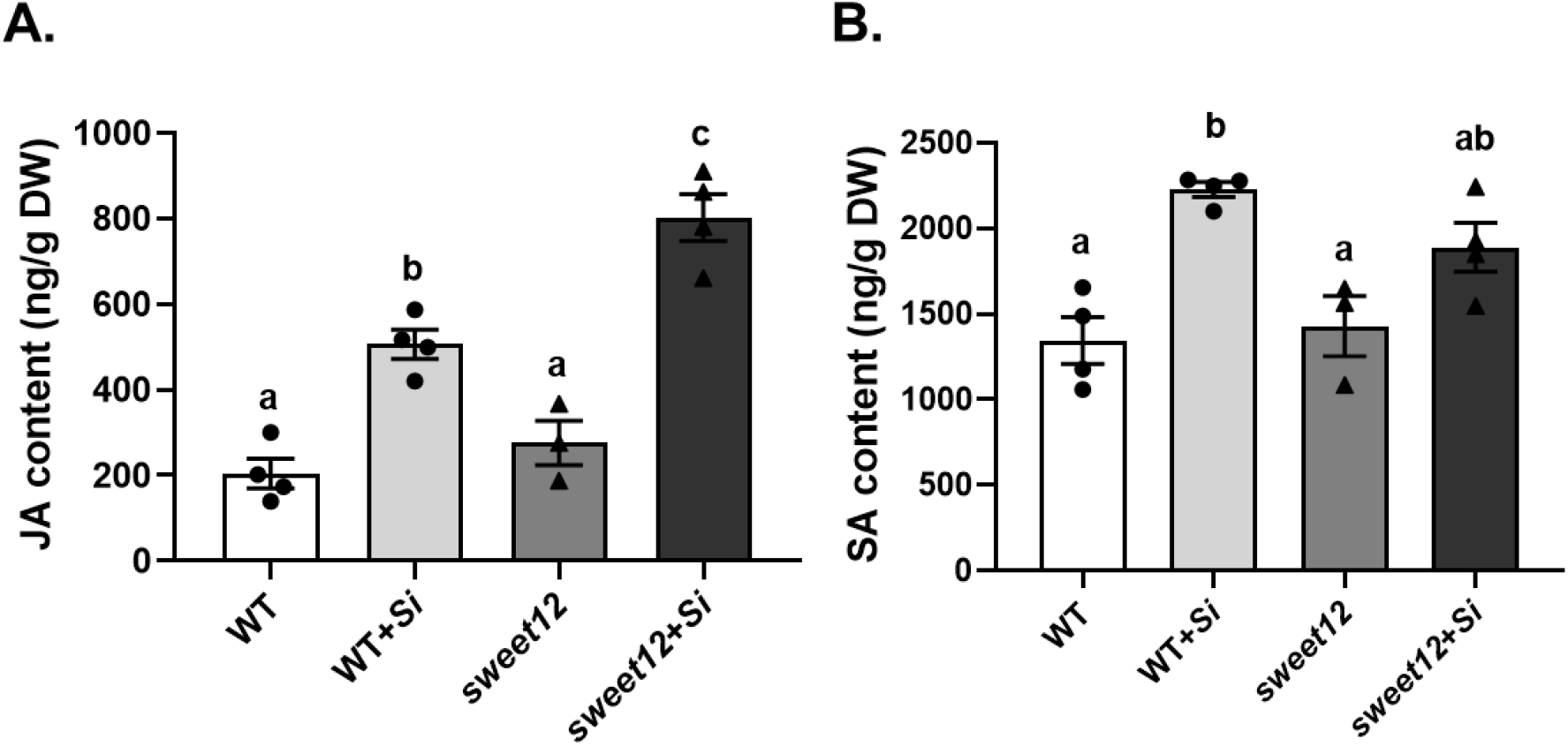
Defense phytohormones after *S. indica-*colonization. Levels of **A**. Jasmonic Acid (JA) **B**. Salicylic acid (SA) at 7 days of *S. indica* inoculation. The seedlings at 7 dpi were harvested, lyophilized, powdered and analysed by using LC/MS. Bars are means ± SEM of at least three independent replicates (n=3*200=600). Different letters indicate significance difference after ANOVA analysis (p < 0.05, one-way ANOVA with Tukey’s test).

### 3.8. Transcriptome analysis shows SWEET12 absence has significant alteration of DEGs during S. indica interaction

We performed transcriptome analysis to gain deeper insights into the global transcriptional responses in *S. indica*-colonized roots (7 dpi) in the absence of SWEET12. Transcriptome analysis revealed that the absence of SWEET12 causes extensive transcriptional reprogramming during *S. indica* colonization. At 7 dpi, WT roots exhibited 1,315 up- and 373 downregulated DEGs, whereas *sweet12* roots showed 2,480 up- and 1,121 downregulated DEGs (**Fig. 8A; Supplementary Data Sets 1–2**). DiVenn analysis demonstrated 672 DEGs unique to WT, 2,541 unique to *sweet12* with *S. indica*, and 1,006 unique to *sweet12* controls (**Fig. 8B**). A total of 747 DEGs were commonly upregulated in both WT and *sweet12* upon colonization, while 568 and 1,726 were uniquely induced in WT and s*weet12*, respectively; conversely, 282 and 1,037 DEGs were uniquely downregulated, with only 84 shared (**Fig. 8C**). Notably, seven genes including *CARBONIC ANHYDRASE 2* (*CA2*), Photosystem II reaction center protein D (*PSBD*), a Glycine-rich protein family protein (*AT2G05540*), *SALICYLIC ACID-BINDING PROTEIN 3* (*SABP3*), *SENESCENCE ASSOCIATED GENE 1* (*SAG1*), a Pollen allergen and extensin family gene (*AT2G27385*), and *Cold Shock Protein 4* (*CSP4*) showed opposite regulation between genotypes, being upregulated in WT but downregulated in *sweet12* (**Fig. 8C**). GO and network analyses revealed that WT roots displayed stronger enrichment of DEGs in external stimulus responses, photosynthesis, carbohydrate metabolic process, while *sweet12* roots showed no or reduced enrichment of these processes and induction of organelle organization, cell cycle, reproductive process pathways and repression of amide and peptide pathways (**Fig. 8D; Fig. S2A–C**). Clustering and cluster wise GO analyses further highlighted that WT colonization induced genes linked to stress response, ion transport, secondary metabolism, and interspecies interactions (clusters 2, 3, 4 and 6; **Fig. 9A; Supplementary Data Set 3**), whereas *sweet12* roots showed weaker induction of these processes but unique activation of peptide metabolism, translation, defense response, and salicylic acid signaling, alongside strong repression of carbohydrate, lipid, ion, and secondary metabolic pathways. Comparative analysis also showed that in WT roots, *S. indica* colonization upregulated *SWEET12, SUS3, CINV6*, and invertases important for sugar mobilization, whereas *sweet12* roots induced alternative transporters such as *STP4, STP14, SWEET8, and SWEET15* (**Fig. 9D; Fig. S2A**), consistent with altered sugar levels and enhanced colonization. Nutrient transport was also differentially regulated, with *sweet12* roots displaying increased expression of nitrate transporters (*NRT1*.*5, NRT2*.*1, NRT2*.*4*) but reduced expression of iron, potassium, and phosphate transport genes crucial for symbiosis and growth promotion response (**Fig. S3B**). Moreover, defense-related SA regulated pathways were strongly suppressed in *sweet12*, with reduced expression of cytochrome P450s, indole glucosinolate biosynthesis genes, chitinases, SA pathway components, and transcription factors including *WRKY33, WRKY70*, while WT roots showed stronger induction of glutathione S-transferases (**Fig. S4A–B**). Cluster comparisons (**Fig. 9A**) showed that although similar categories of biological processes were enriched in both control WT and *sweet12* roots, the extent and direction of their regulation differed, with *sweet12* exhibiting a skewed transcriptional response compared to WT

**Fig. 8.**
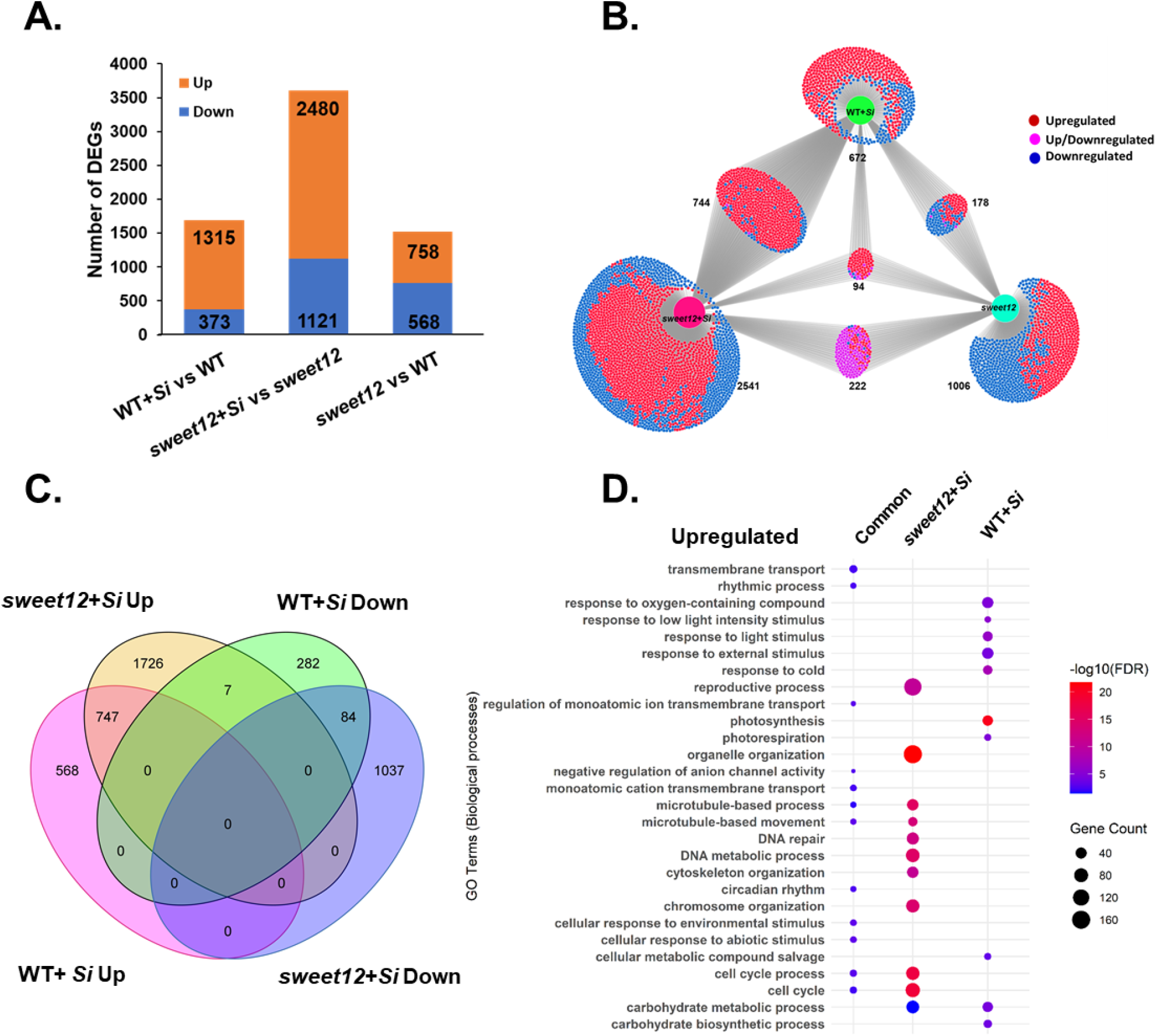
Transcriptomics analysis of *sweet12* control and *S. indica-*colonized *sweet12* mutant roots compared to WT. **A**. Numbers of differentially expressed genes (DEGs) in WT control, WT with *S. indica, sweet12* mutant control and *sweet12* mutant roots at 7 dpi (Log2FC = 0.58). **B**. DiVenn diagram showing the numbers of DEGs in WT with *S. indica, sweet12* mutant and *sweet12* with *S. indica*. **C**. Venn diagram showing the numbers of DEGs up- and downregulated in WT with *S. indica*, and *sweet12* with *S. indica* (Log2FC = 0.58). **D**. DEGs belonged to different biological processes upregulated in response to *S. indica* in roots at 7 dpi in WT and *sweet12* roots. The size and color of the circle in the graph represents number of genes and fold enrichment (- Log10FDR), respectively of corresponding biological process.

**Fig. 9.**
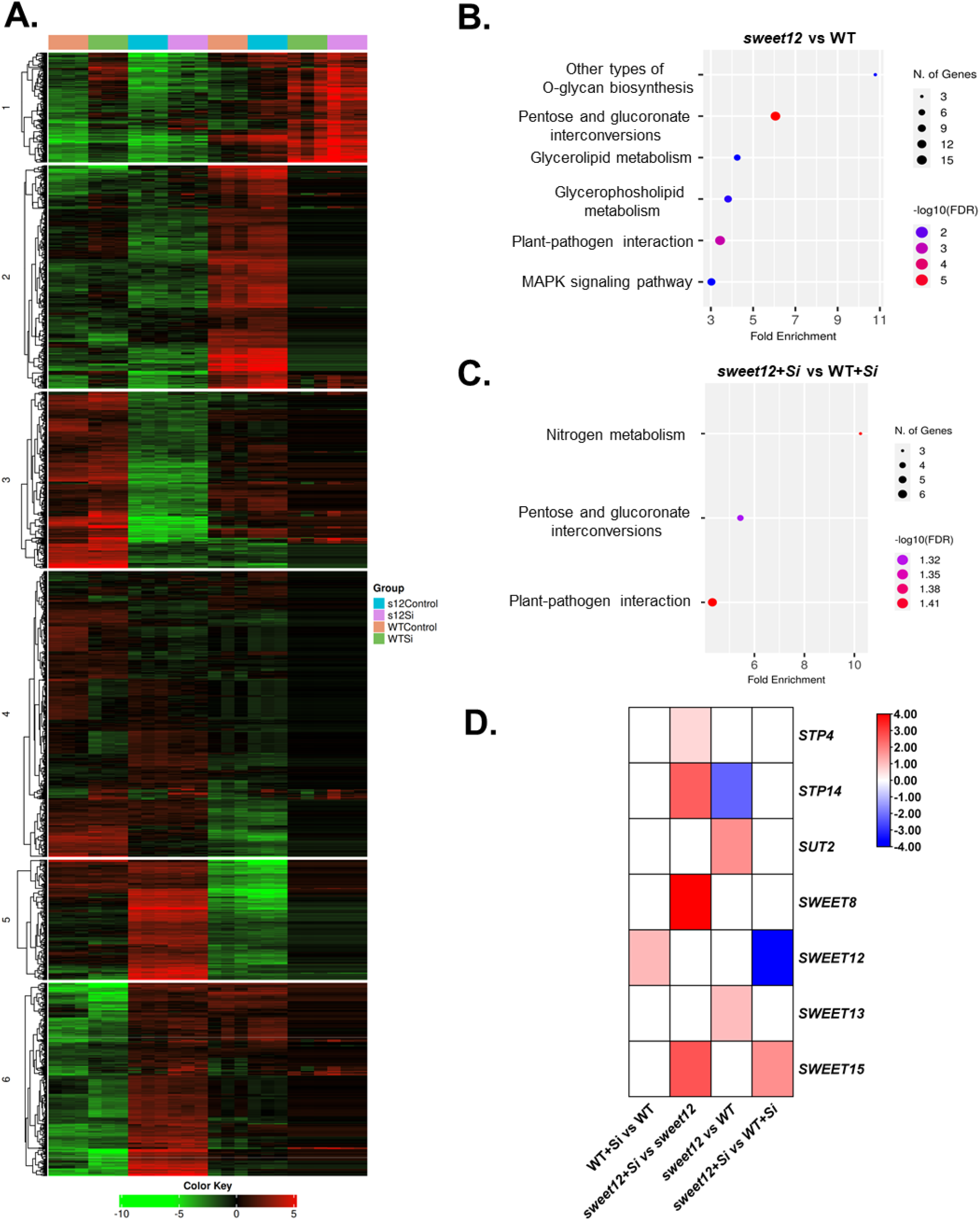
Clustering and KEGG pathway analysis of *S. indica-*colonized WT and *sweet12* mutant roots. **A. Clustering:** It was performed using raw read count data of all samples. All DEGs in *S. indica*-colonized *sweet12* roots and *S. indica*-colonized WT roots were as input in iDEP tool and k-Means clustering analysis was performed at -Log10FDR = 0.05. **B**. KEGG enrichment analysis of upregulated DEGs in *S. indica*-colonized WT roots over its non-colonized control. **C**. KEGG enrichment analysis of upregulated DEGs in *S. indica*-colonized *sweet12* roots over its non-colonized control. The size and color of the circle in the graph represents number of genes enriched and fold enrichment (-Log10FDR = 0.05), respectively of a particular corresponding KEGG pathway. **D. Heat map of differentially expressed sugar transporters**. Log2FC of DEGs were used to generate heat map of Sugar transporters.

## 4. Discussion

In terrestrial plants, carbon exchange is mediated by the induction of specific sugar transporters. Modulation of these transporters allows microbes to redirect sugar flux into the apoplast, where they can access and utilize these nutrients. In symbiotic relationships, the microbial partner primarily acquires carbon in the form of sugars from the host plant (Hawkins et al. 2023). Genome analysis in *Serendipita indica* revealed 19 putative hexose transporters (HXTs), with HXT5 being strongly induced during root colonization, coinciding with maximal fungal growth inside roots. *Si*HXT5 is s a high-affinity, H^+^-dependent sugar transporter, capable of importing multiple monosaccharides with a preference for glucose (Rani et al. 2015, Raj et al. 2021). However, the specific plant sugar transporters responsible for delivering sugars to *S. indica* remain unidentified. The mutualistic interaction between *S. indica* and Arabidopsis enhances sugar availability through the activation of invertases and sucrose synthases in the plant, which is essential for promoting growth and enhancing defense responses (Opitz et al. 2021). Identifying the host sugar transporters that play a critical role in supplying sugars to *S. indica* and maintaining sugar homeostasis is vital for understanding the sustainability of this symbiosis.

Our study highlights the central role of the sugar transporter SWEET12 in regulating Arabidopsis–*Serendipita indica* symbiosis. We show that SWEET12 is rapidly induced by both fungal colonization and the fungal elicitor cellotriose (CT), consistent with earlier reports of SWEET induction during microbial interactions (Pérez-Alonso et al. 2022, Gandhi et al. 2024). The tissue-specific activation of SWEET12 in roots, particularly in the cortex, vasculature, and root hairs, suggests that this transporter may function at multiple entry points of carbon flux to the apoplast, thereby providing the fungus with access to host-derived sugars, in a spatially controlled manner. This is reminiscent of other symbiotic systems, where root-localized SWEETs contribute to sugar secretion during arbuscular mycorrhizal colonization (Manck-Götzenberger and Requena, 2016). Loss-of-function of SWEET12 led to growth inhibition instead of growth promotion, as expected from mutation of a sugar transporter, where sugar availability is reduced. However, the observation that *sweet12* mutants exhibit increased *S. indica* colonization, despite reduced growth promotion appeared contradictory. However, several mechanisms can explain this outcome based on our whole genome transcriptome. In the absence of SWEET12, soluble sugars such as sucrose, glucose, and fructose accumulate intracellularly rather than being exported from source (leaf) to sink (root) in a regulated manner, which may alter osmotic balance, weaken cell wall integrity, or trigger compensatory activity of alternative transporters such as STP4, STP14, SWEET8, and SWEET15, resulting in exploitation by the fungus. Consequently, the fungus colonizes roots more aggressively to derive more sugars from the minimal pool, as also evident from enhanced hyphal penetration in *sweet12* roots, but without conferring the balanced growth benefits observed in wild - type plants. This finding supports the idea that controlled sugar efflux is essential for maintaining mutualism. Thus, SWEET12 might act as a “carbon valve” releasing sugars in a regulated manner that promotes beneficial colonization while avoiding uncontrolled fungal expansion. Excess colonization without growth benefit in *sweet12* mutant parallels observations in other systems where deregulated sugar allocation enhances pathogen proliferation (Chen et al. 2010, Gebauer et al. 2017). In Arabidopsis, *S. indica* over-colonization with no growth promotion phenotype has been also reported in various mutants affecting multiple signaling pathways downstream such as *mdar2/mdar5, pen2, etr1, ein2, ein3, cyp79b2*/*cyp79b3, myb34/51/122, pad3, and cngc19* (Vadassery et al. 2009b, Jacobs et al. 2011, Camehl et al. 2010, Nongbri et al. 2012, Lahrmann et al. 2015, Jogawat et al., 2020), whereas Arabidopsis mutants such as *cbl7, wrky6, adc1*, and *adc2* show no growth promotion as a result of less colonization (Bakshi et al. 2015, Kundu et al. 2022, Pérez-Alonso et al. 2022). In a study *S. indica* has been shown to alter the expression of genes involved in sugar metabolism, such as invertases and sucrose synthases. Mutants of these genes (*sus1/2/3/4* and *cinv1/2*) exhibit altered sugar pools and the *cinv1/2* mutant displayed higher *S. indica* colonization compared to WT roots (Opitz et al. 2021).

SWEET12 also influences defense phytohormone signaling. We observed that JA levels were elevated in *sweet12* mutants upon *S. indica* colonization, whereas SA accumulation and SA-associated markers were reduced. This might co-relate with increased colonization and plants trying to activate MAMP triggered immunity. CNGC19 is an important Ca^2+^ channel that maintains this robust innate immunity and is essential for maintaining balanced symbiosis (Jogawat et al. 2020). Loss of CNGC19 leads to excessive fungal colonization and growth inhibition like in *sweet12* mutant due to impaired Ca^2+^ signaling, compromised MAMP-triggered immunity, reduced jasmonate accumulation, and failure to regulate indole glucosinolates (Jogawat et al. 2020). Further, given the established link between soluble sugars, SA signaling, and defense gene activation (Gebauer et al. 2017, Yamada and Mine 2024), SWEET12 likely integrates carbon allocation with immune modulation. This is further supported by transcriptome data showing attenuated induction of SA-responsive genes, WRKY transcription factors, and glucosinolate biosynthetic genes in *sweet12* roots. Together, these findings suggest that SWEET12 helps maintain the defense–growth tradeoff, enabling plants to reap the benefits of symbiosis while preventing excessive fungal colonization. Various studies have already shown the involvement of secondary metabolites and their biosynthesis pathway genes in *S. indica*-symbiosis (Camehl et al. 2013, Nongbri et al. 2012, Lahrmann et al. 2015). In *sweet12* mutant roots, we observed reduced expression of these genes which includes *BCAT7, APK1, ASA1, CYP79F1, CYP81F2, IGMT1, IGMT2, MYB29, MYB51, MYB122, NIT1, NIT2, NIT4, PAD3*, and *WRKY33*. These genes are shown to restrict *S. indica* colonization level to maintain growth promotion activity. For instance, double mutant *cyp79b2/cyp79b3* and *pad3* mutant show reduced indole-3-acetaldoxime (IAOx)-derived defense metabolites which results in increased colonization with loss of growth promotion activity (Nongbri et al. 2012). *AtSWEET12* is induced by *P. syringae* infection in leaf and acts as controller of sucrose transport together with SWEET11 during foliar infection. In the absence of *sweet12*, there is no heterodimer formation by SWEET11 and SWEET12 which leads to over loading of sucrose into apoplast which subsequently leads to increased load of bacterial cells (Fatima and Senthil-Kumar 2021). This corroborates our findings in Arabidopsis roots during *S. indica* infection with respect to higher level of different sugars, over-colonization and growth inhibition upon *S. indica* colonization. Transcriptome reprogramming in the absence of SWEET12 underscores its broader regulatory role. Compared to WT, *sweet12* mutants displayed enhanced induction of peptide metabolism and translation but repression of carbohydrate, lipid, ion, and secondary metabolic pathways. Particularly striking is the reduced expression of nutrient transporters for iron and potassium, both of which are known to modulate mycorrhizal and root endophyte interactions (Ito et al. 2024, Pérez-Alonso et al. 2022). This suggests that SWEET12 indirectly supports nutrient homeostasis during symbiosis, likely by ensuring an appropriate sugar–nutrient exchange ratio.

Collectively, these findings suggest that SWEET12 functions as a “carbon valve” releasing sugars in a regulated manner that supports beneficial colonization while preventing uncontrolled fungal proliferation. By maintaining this balance, SWEET12 emerges as an essential component of the mutually beneficial relationship between Arabidopsis and *S. indica*, underscoring its role in fungal colonization, defense regulation, and plant growth promotion. In addition, SWEET12 acts as a key regulator of immune competence and the carbon–defense balance, ensuring controlled nutrient allocation to *S. indica* while simultaneously restricting over-colonization. In its absence, this balance collapses, shifting the interaction toward a parasitic-like state in which excessive fungal growth occurs at the expense of host fitness (**Fig. 10A–B**).

**Fig.10.**
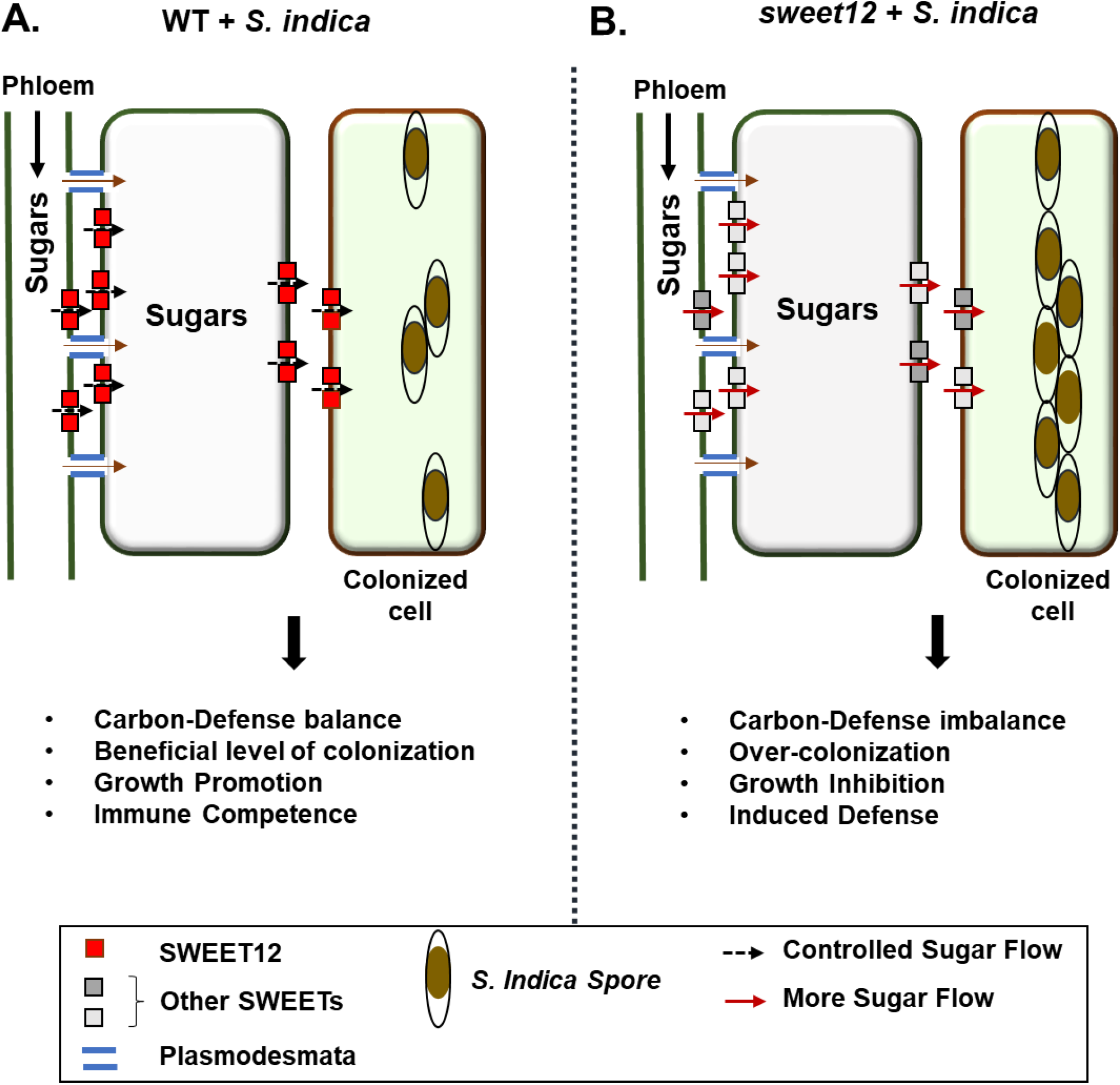
Proposed functional role of Arabidopsis SWEET12 during S. *indica* symbiosis. **A**. In WT plants upon *S. indica* colonization, sugar synthesis and mobilization is induced. In this symbiosis, SWEET12 functions like a “carbon valve” mobilizing sugars in a controlled manner that sustains beneficial colonization while avoiding non-beneficial fungal overload. By maintaining this balance, SWEET12 appears as an essential component of the mutually beneficial relationship between Arabidopsis and *S. indica*, underscoring its role in fungal colonization, defense regulation, and growth promotion. In addition, SWEET12 acts as a key regulator of immune competence and the carbon–defense balance, ensuring controlled nutrient allocation to *S. indica* while simultaneously restricting over-colonization. **B**. In *sweet12* mutant, this balance collapses, shifting the interaction toward a pathogen-like state in which excessive fungal growth occurs to extract more sugars, at the expense of host fitness. Here, the single line arrows indicate flow of sugar molecules.

## Data Availability

The data that support the findings of this study are available from the corresponding author upon reasonable request.

## Acknowledgements and funding

AJ acknowledges Department of Biotechnology, Government of India for providing fellowship and research funding through MK Bhan-Young Researcher Fellowship (File No. HRD-16016/2/2023-AFS-DBT) program. MS and SHM were funded by JRF from MK Bhan-Young Researcher Fellowship of AJ. AMN acknowledges DST-INSPIRE fellowship. We acknowledge NIPGR for core grant, Central instrumentation Facility, Metabolome Facility. We thank Senthil-Kumar Muthappa (NIPGR) for providing mutant of *SWEET12*. We thank Cécile Vriet (University of Poitiers) for gifting GUS line of *SWEET12*. We also acknowledge Neuberg Supratech Laboratories, Ahmadabad, India for RNA sequencing.

## Authors’ contributions

AJ and JV designed the experiments, analyzed the results and wrote the manuscript. AJ, MS, SHM and AMN performed the experiments. DG performed GS-MS and LC-MS experiments. All authors approved the manuscript.

## Supplementary Table

**Supplementary Table S1**. List of primer pairs used in this study.

## Supplementary Figures

**Fig. S1**. RNA genotyping of *sweet12* mutant line (SALK_031696).

**Fig. S2**. Network analysis of DEGs.

**Fig. S3**. Transcripts level of sugar and nutrient transport related genes in Arabidopsis wild-type (WT) and *sweet12* mutant 7 dpi in roots upon *S. indica* colonization.

**Fig. S4**. Transcripts level of defense related genes in Arabidopsis wild-type (WT) and *sweet12* mutant 7 dpi in roots upon *S. indica* colonization.

## Supplementary Datasets

**Supplementary Dataset 1**. List of DEGs up- and downregulated in the RNA sequencing of wild type + *S. indica* with functional annotation.

**Supplementary Dataset 2**. List of DEGs up- and downregulated in the RNA sequencing of *sweet12* + *S. indica* with functional annotation.

**Supplementary Dataset 3**. List of genes, biological processes and KEGG pathways enriched in clustering analysis of different groups.

