## Supplementary Information for "Arabidopsis SWEET12 regulates sugar allocation and defense responses to sustain beneficial association with *Serendipita indica* in roots"

**Supplementary Table 1: List of primer pairs used in this study**

| S.N. | Primer name | Sequence | Experiment |
|---|---|---|---|
| 1 | SWEET1RTF | TGTGGTACTTCGTGGTTCGT | RT-PCR |
| 2 | SWEET1RTR | TTGCAGTGTCCCTAATGCAC | RT-PCR |
| 3 | SWEET2RTF | TGGTGTTTGCTGTGGTTGGA | RT-PCR |
| 4 | SWEET2RTR | CGGCGAGGCAAACATTGAAA | RT-PCR |
| 5 | SWEET3RTF | GTCGGCATCCTTCTCGAATCT | RT-PCR |
| 6 | SWEET3RTR | CTGTCGTTAAGCCGAACCCA | RT-PCR |
| 7 | SWEET4RTF | CATTGCCGGCATTTGTGGAA | RT-PCR |
| 8 | SWEET4RTR | TGCGCAATTTAGAACCGTGG | RT-PCR |
| 9 | SWEET5RTF | ATGGCGGTGGTGATCTTCTG | RT-PCR |
| 10 | SWEET5RTR | CATGACGGTGAGAGGAGCAG | RT-PCR |
| 11 | SWEET6RTF | ATGGTGCATGAACAGTTGAA | RT-PCR |
| 12 | SWEET6RTR | CAACCGATGTTGTTGACGACC | RT-PCR |
| 13 | SWEET7RTF | TTACGGACTACCAACGGTGC | RT-PCR |
| 14 | SWEET7RTR | TTTGGCGGCCACAATAAACG | RT-PCR |
| 15 | SWEET8RTF | CTTTGGTCTCTTTGCCGCAC | RT-PCR |
| 16 | SWEET8RTR | TGAACCACAGGGAGACCGTA | RT-PCR |
| 17 | SWEET9RTF | CCACGAAAACGGATTTGCCA | RT-PCR |
| 18 | SWEET9RTR | GCCTCACCGATCCTTCAACA | RT-PCR |
| 19 | SWEET10RTF | CTTTTTCGTCTGCCTTGCCC | RT-PCR |
| 20 | SWEET10RTR | ATCCATAGCATCGCGCTGAA | RT-PCR |
| 21 | SWEET11RTF | GGCACAGTTTCATCCCCTGA | RT-PCR |
| 22 | SWEET11RTR | GGAAGAGGACTGCTTGCCAT | RT-PCR |
| 23 | SWEET12RTF | TCTGTCTGCGTTTTTGCTGC | RT-PCR |
| 24 | SWEET12RTR | GAGCCATATGACCGCACTGA | RT-PCR |
| 25 | SWEET13RTF | CGTCAGTGTTTTCGCAGCTC | RT-PCR |
| 26 | SWEET13RTR | GTAGAAGAGCCACGTGACGG | RT-PCR |
| 27 | SWEET14RTF | ACTTCTACGTTGCGCTTCCA | RT-PCR |
| 28 | SWEET14RTR | ACCGTTTTGGGCTTCTCTGT | RT-PCR |
| 29 | SWEET15RTF | TCCTCCCCTCCAAGTCTCTG | RT-PCR |
| 30 | SWEET15RTR | AGCGTGAAGGGCATGTACTC | RT-PCR |

| | | | |
|---|---|---|---|
| 31 | SWEET16RTF | GCGATTGCGGGAACAAGAAC | RT-PCR |
| 32 | SWEET16RTR | CGTCACAACCGTTTTAATAGCTGA | RT-PCR |
| 33 | SWEET17RTF | ATTACGGCATCGTCACTCCC | RT-PCR |
| 34 | SWEET17RTR | TCGCTTCCACATCCACTGTT | RT-PCR |
| 35 | AtACTIN2RTF | TCAGATGCCCAGAAGTCTTGTTC | RT-PCR |
| 36 | AtACTIN2RTR | GTGGATTCCAGCAGCTTCCA | RT-PCR |
| 37 | SiTEF1RTF | TCGTCGCTGTCAACAAGATG | RT-PCR |
| 38 | SiTEF1RTR | GAGGGCTCGAGCATGTTGT | RT-PCR |
| 39 | SWEET12RGF | TCTGTCTGCGTTTTTGCTGC | RNA Genotyping |
| 40 | SWEET12RGR | GAGCCATATGACCGCACTGA | RNA Genotyping |

**Supplementary Figures:**

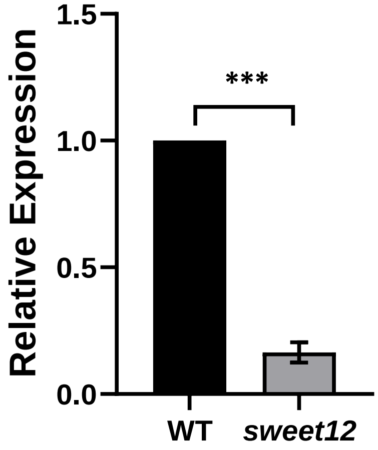


**Fig. S1. RNA genotyping of *sweet12* mutant line (SALK_031696).** RNA was extracted, cDNA was prepared and *SWEET12* transcript level was checked by using *SWEET12* real time-PCR primers in 10 days old seedlings. Data represents mean fold change ± SEM of n = 3 (3*10=30). Asterisks indicate significant difference after Student's *t*-test, $p < 0.0001$.

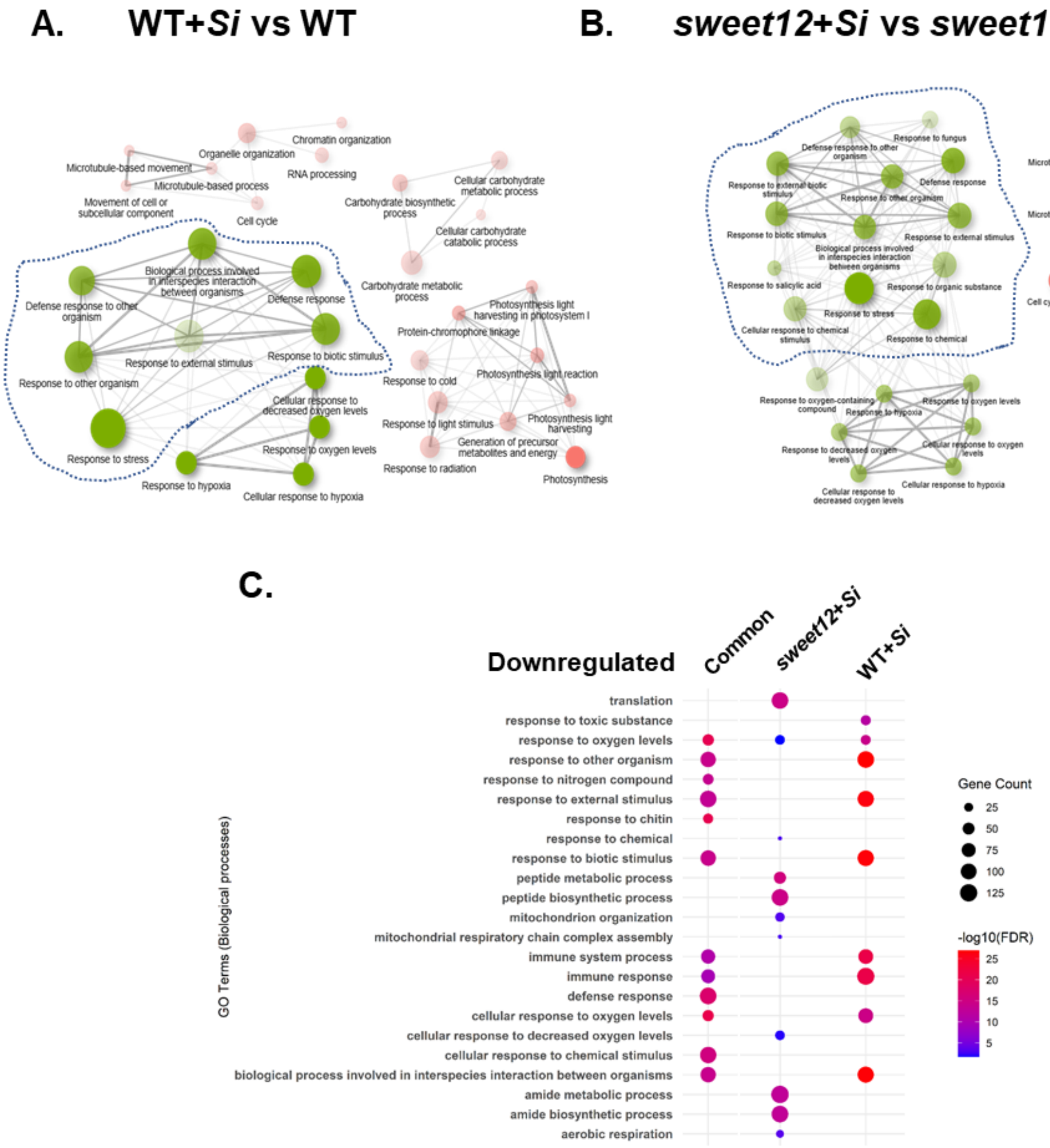


**Fig. S2. Network analysis of DEGs. A. *S. indica*-colonized WT roots B. *S. indica-sweet12* mutant roots.** Based on enrichment of biological processes, network of associated DEGs was generated. In the network, two pathways (nodes) are likely to be linked if ≥30% genes are shared. Green and red color represents up- and down-regulated pathways, respectively. Color intensity suggests comparative significant enrichment of gene sets whereas size indicates number of genes in the pathway. Thickness of edges represents relative number of overlapped genes. The pathways are sorted based on a FDR cutoff (0.05) and 1.5-fold change. Only the top 20 pathways are shown. **C.** DEGs belonged to different bioprocesses downregulated in response to *S. indica* in roots at 7 dpi. The size and color of the circle in the graph represents number of genes and fold enrichment (-Log10FDR), respectively of corresponding biological process.

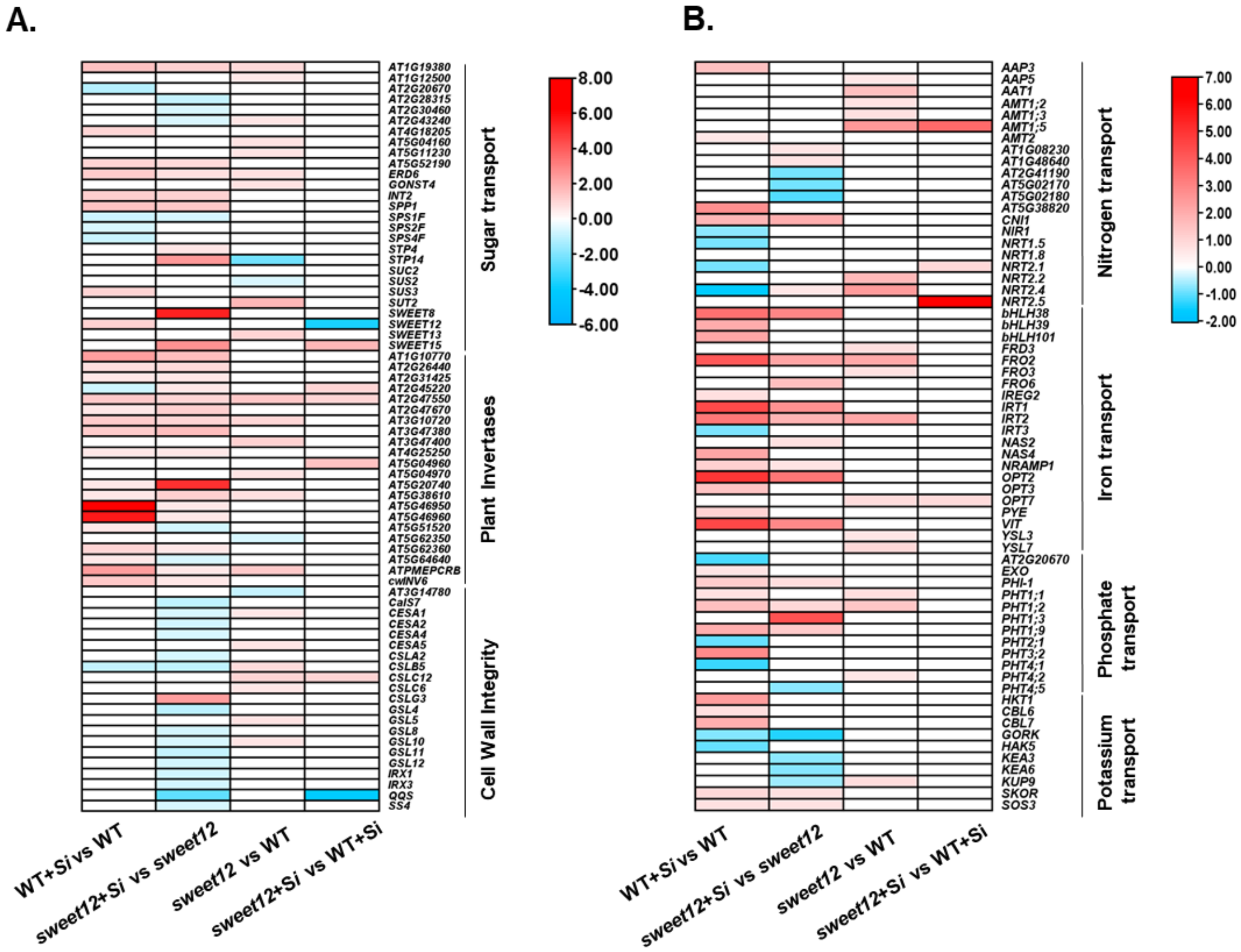


**Fig. S3. Transcripts level of sugar and nutrient transport related genes in Arabidopsis wild-type (WT) and *sweet12* mutant 7 dpi in roots upon *S. indica* colonization. A.** Heat map of sugar transport, plant sugar invertases and cell wall integrity genes **B.** Heat map of nitrogen transport, iron transport, phosphate transport and potassium transport. Differential expression was calculated over non-colonized control and the log2fold-change change was taken at cutoff of 0.58.

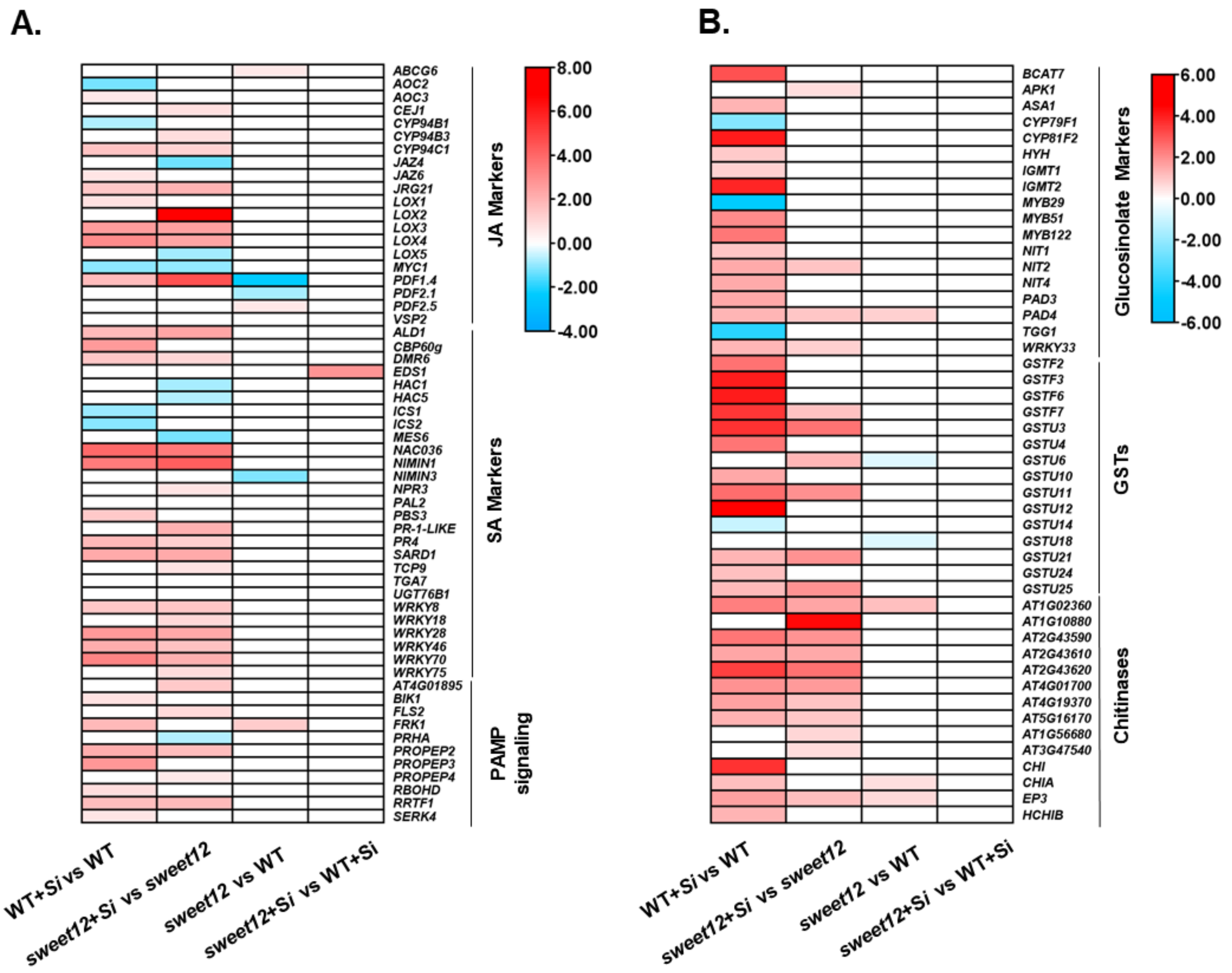


**Fig. S4. Transcripts level of defense marker genes in Arabidopsis wild-type (WT) and *sweet12* mutant 7 dpi in roots upon *S. indica* colonization. A.** Heat map of JA marker, SA marker, and plant pathogen response genes. **B.** Heat map of glucosinolate pathway genes, GSTs, and chitinase genes. Differential expression was calculated over non-colonized control and the log2fold-change change was taken at cutoff of 0.58.
